# Transsynaptic mapping of *Drosophila* mushroom body output neurons

**DOI:** 10.1101/2020.09.22.309021

**Authors:** Kristin M Scaplen, Mustafa Talay, John D Fisher, Raphael Cohn, Altar Sorkaç, Yoshinori Aso, Gilad Barnea, Karla R Kaun

## Abstract

The Mushroom Body (MB) is a well-characterized associative memory structure within the *Drosophila* brain. Although previous studies have analyzed MB connectivity and provided a map of inputs and outputs, a detailed map of the downstream targets is missing. Using the genetic anterograde transsynaptic tracing tool, *trans-*Tango, we identified divergent projections across the brain and convergent downstream targets of the MB output neurons (MBONs). Our analysis revealed at least three separate targets that receive convergent input from MBONs: other MBONs, the fan shaped body (FSB), and the lateral accessory lobe (LAL). We describe, both anatomically and functionally, a multilayer circuit in which inhibitory and excitatory MBONs converge on the same genetic subset of FSB and LAL neurons. This circuit architecture provides an opportunity for the brain to update information and integrate it with previous experience before executing appropriate behavioral responses.

**Highlights:** -The postsynaptic connections of the output neurons of the mushroom body, a structure that integrates environmental cues with associated valence, are mapped using *trans*-Tango.

-Mushroom body circuits are highly interconnected with several points of convergence among mushroom body output neurons (MBONs).

-The postsynaptic partners of MBONs have divergent projections across the brain and convergent projections to select target neuropils outside the mushroom body important for multimodal integration.

-Functional connectivity suggests the presence of multisynaptic pathways that have several layers of integration prior to initiation of an output response.

## Introduction

The brain comprises intricate neural networks in which information iteratively converges and diverges to support learning, memory, and behavioral flexibility. Knowledge of the neural connectivity that underlies these networks is essential to our understanding of how the brain functions. *Drosophila melanogaster* provides a powerful opportunity to map the fundamental architecture of neural circuits due to its complex yet tractable brain. With a nervous system of approximately 100,000 neurons and a rich genetic toolkit that offers the potential to selectively manipulate subsets of neurons in behaving animals, significant effort has been devoted to establishing a detailed map of neural connectivity in the fly (Aso et al., 2014a; Bates et al., 2020; Couto et al., 2005; Deng et al., 2019; Eichler et al., 2017; Eschbach et al., 2020; Fishilevich and Vosshall, 2005; Frechter et al., 2019; Grabe et al., 2015; Kondo et al., 2020; Li et al., 2020; Marin et al., 2020; Otto et al., 2020; Peng et al., 2011; Shao et al., 2014; Takemura et al., 2017; Zheng et al., 2018).

The mushroom body (MB) is a prominent neuropil structure that receives input from multiple sensory modalities (Ehmer and Gronenberg, 2002; Li and Strausfeld, 1999; Liu et al., 2006; Liu et al., 2016; Marin et al., 2020; Marin et al., 2002; Strausfeld and Li, 1999a b;, Vogt et al., 2016; Vogt et al., 2014; Wang et al., 2016; Yagi et al., 2016; Zars, 2000) and has a well-established role in olfactory learning and memory (Davis, 1993; de Belle and Heisenberg, 1994; Heisenberg, 1998, 2003; Heisenberg et al., 1985; Pascual and Preat, 2001; Zars et al., 2000). It comprises thousands of densely packed parallel axonal fibers of Kenyon cells that are organized into three separate lobes (α/β, α′/β′, γ) (Crittenden et al., 1998; Ito et al., 1997; Ito et al., 1998; Strausfeld, 2002; Strausfeld and Li, 1999a, b). Kenyon cells that are intrinsic to the MB receive divergent input from diverse combinations of olfactory glomeruli (Caron et al., 2013; Gruntman and Turner, 2013). They also receive organized valence-related input from dopamine neurons (DANs) and converge onto 34 different MB output neurons (MBONs) (Aso et al., 2014a; Eichler et al., 2017; Eschbach et al., 2020; Takemura et al., 2017). This architecture positions the MB as a high-level integration center for the representations of olfactory cues and their perceived valence.

With the development of split-Gal4 lines that provide selective genetic access to precise neuronal populations (Aso et al., 2014a), detailed patterns of MB neural circuits have emerged over the past decade describing a compartmentalization of the MB lobes by arborization patterns of innervating dopamine neurons (DANs) and MBONs (Aso et al., 2014a; Eichler et al., 2017; Eschbach et al., 2020; Takemura et al., 2017). In addition to a detailed description of the input and output innervation patterns of the MB, projection patterns of MBONs were characterized, and several neuropil structures were identified as sites of convergence for the MBONs, including the lateral horn (LH), crepine (CRE), superior medial (SMP), intermediate (SIP), and lateral (SLP) protocerebrum (Aso et al., 2014a). Within these convergent neuropil structures, MBON axons were proposed to synapse onto axons of other MBONs, and dendrites of DANs, interneurons and projection neurons. These convergent neuropil structures, however, are characterized by highly complex arborizations of dendrites and axons making it challenging to identify the specific neural components that receive synaptic input from various MBONs.

Identification of postsynaptic partners of specific neurons was previously limited to using computational approaches that identify potential candidate neurons based on their overlapping arborization patterns or using tools such as fluorescent protein reconstitution across synaptic partners to confirm connectivity (Chiang et al., 2011; Feinberg et al., 2008; Jefferis et al., 2007; Li et al., 2016; Lin et al., 2013; Macpherson et al., 2015; Shearin et al., 2018; Wolff et al., 2015). More recently, the *Drosophila* field has made a concerted effort to map synaptic connections across the fly brain using whole brain electron microscopy (EM) data (Li et al., 2020; Ohyama et al., 2015; Schneider-Mizell et al., 2016; Xu et al., 2020; Zheng et al., 2017; Zheng et al., 2018). Although EM data offers synaptic resolution, it is labor intensive and does not account for potential variability in synaptic connectivity that exists across animals. We sought to complement the EM anatomic data by mapping the postsynaptic partners of all MBONs using the genetic anterograde transsynaptic tracing tool, *trans*-Tango (Talay et al., 2017). We found that MB efferent pathways are highly interconnected, including several points of convergence among MBONs. We also revealed direct connections between the MBONs and two additional regions, the fan shaped body (FSB) and the lateral accessory lobe (LAL). We further describe, both anatomically and functionally, a multilayer circuit that includes GABAergic and cholinergic MBONs that converge on the same subset of FSB and LAL postsynaptic neurons, providing an opportunity to integrate information processing before executing behavior. We propose that multilevel integration across brain regions is critical for updating information processing and memory.

## Results

### Divergence and convergence of the MBONs circuits

Circuit convergence, divergence, and re-convergence can be found throughout the nervous systems of both invertebrates and vertebrates and plays a pivotal role in providing behavioral flexibility (Eschbach et al., 2020; Jeanne and Wilson, 2015; Man et al., 2013; Miroschnikow et al., 2018; Misic et al., 2014; Ohyama et al., 2015). Given the importance of the MBONs in driving behavioral choice, we first sought to reveal patterns of divergence and convergence by identifying the postsynaptic connections of the MBONs innervating each of the 15 MB compartments using *trans-*Tango. Since *trans*-Tango signals depend on intensity and specificity of split-GAL4 drivers, we selected 28 previously published MBON split-GAL4 lines specific to individual MBONs, or sparse but overlapping subsets of MBONs (Aso et al., 2014a). We combined *trans*-Tango with chemogenetic active zone marker using the *brp-SNAP* knock-in to increase uniformity of neuropil labeling (Kohl et al., 2014).

We successfully identified the postsynaptic connections of 25 split-GAL4 lines (Aso et al., 2014a). *trans-*Tango signals from MB112C (MBON γ1pedc>α/β) and G0239 (MBON α3) were too weak and were excluded from further analysis. In contrast, signals from MB242A (MBON calyx) proved to be too noisy to confidently identify postsynaptic connections. We also employed three new split-GAL4 lines that had more specific expression for γ5β′2a, β′2mp, and α2sc MBONs. Postsynaptic connections of glutamatergic, GABAergic, and cholinergic MBONs vary with regard to the divergence and breadth of their postsynaptic connections (Figure 1 Supplementary videos 1-25). For instance, MB011B, which includes glutamatergic MBONs γ5β′2a, β′2mp, and β′2mp-bilateral has extensive connections across the superior protocerebrum (Figure 1A), whereas MB542B, which includes cholinergic MBONs α′1, α2p3p, α′3 has limited connections within the lateral horn (Figure 1N). The breadth of innervation patterns did not seem to correlate with neurotransmitter type or number of MBONs expressing each split-GAL4.

**Figure 1:**
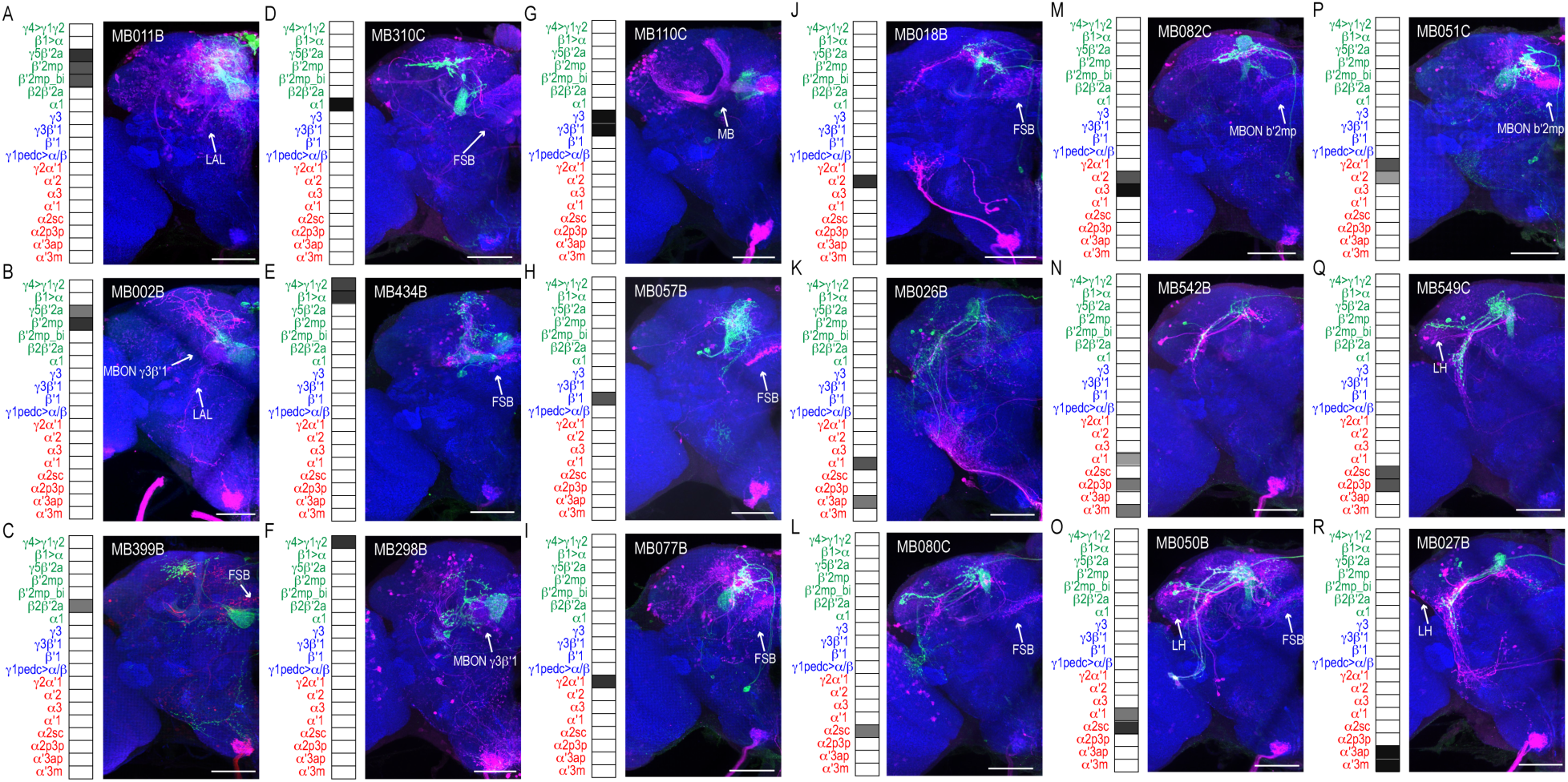
MBONs have divergent connections across the brain. Exemplar max-stacks of glutamatergic MBONs **(A)** MB011B, **(B)** MB002B, **(C)** MB399B, **(D)** MB310C, **(E)** MB434B, **(F)** 298B, GABAergic MBONs **(G)** MB110C and **(H)** MB057B, and cholinergic MBONs **(I)** MB077B, **(J)** MB018B, **(K)** MB026B, **(L)** MB080C, **(M)** MB082C, **(N)** MB542B, **(O)** MB050B, **(P)** MB051C, **(Q)** MB549C and **(R)** MB027B, *trans-*Tango identified postsynaptic connections. For max-stacks: green, presynaptic MBONs, magenta, postsynaptic *trans*-Tango signal, blue, *brp-SNAP* neuropil. A map of the MBONs that are included in the expression pattern in each driver line accompanies each exemplar with the relative expression pattern (greyscale, 1-5) accordingly to FlyLight (https://splitgal4.janelia.org/cgi-bin/splitgal4.cgi). MBON maps are organized by neurotransmitter type: green=glutamatergic, blue=GABAergic, red=cholinergic. Scale bar=50μm.

However, it was clear that some of the data was confounded by split-GAL4 lines that had off-target expression. We excluded off-targeted *trans*-Tango signals by segmenting *trans*-Tango signals that were continuous with MBON terminals (Figure 2A-B) and then quantified the distribution of postsynaptic signals across brain regions in the standard brain (Ito et al., 2014) (Figure 2C-E). Nearly all MBONs have divergent connections across the dorsal brain regions, CRE, SMP, SIP, SLP, LH, as well as FSB, and LAL.

**Figure 2:**
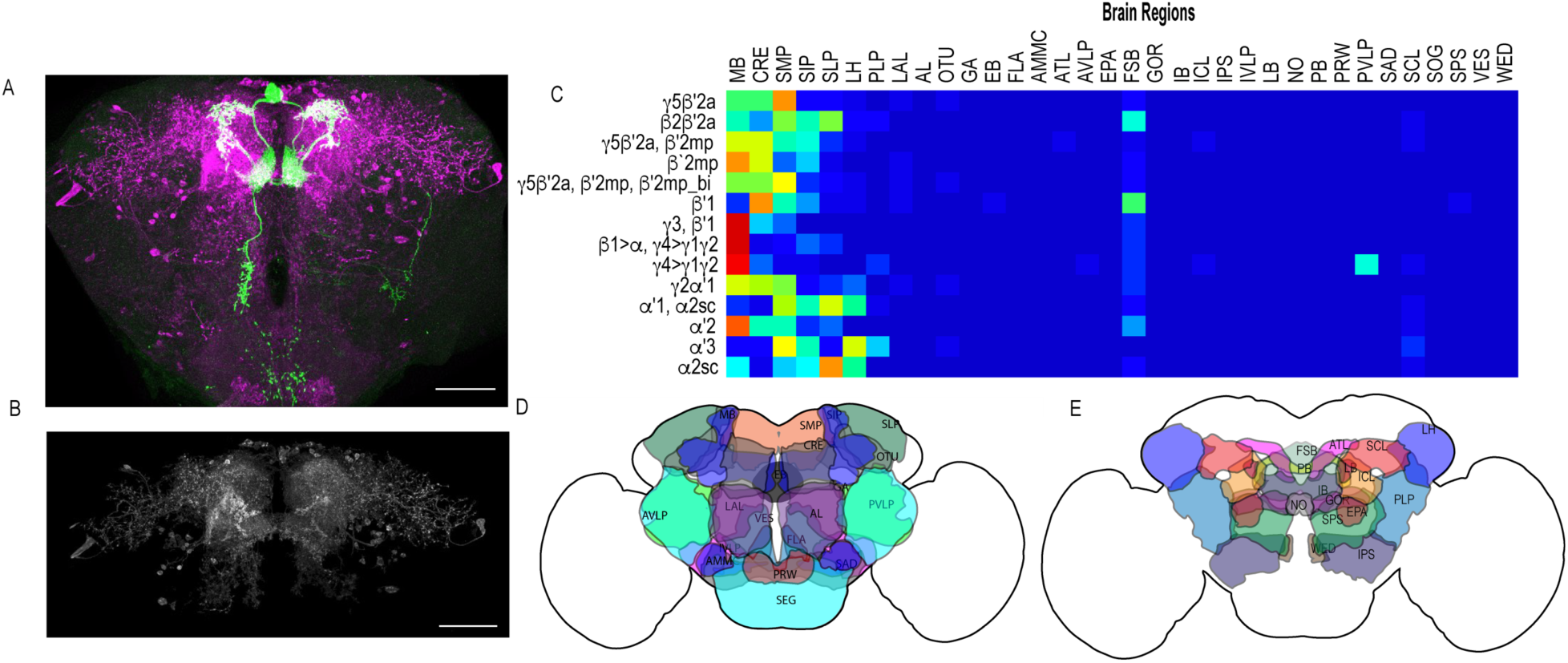
Whole brain distribution of MBON postsynaptic connections overlap. **(A)** Example of presynaptic MBON γ5β′2a (SS01308) and postsynaptic *trans*-Tango signal in a registered brain. For max-stacks: green, presynaptic MBONs, magenta, postsynaptic *trans*-Tango signal. **(B)** Example of segmented *trans*-tango signals that was continuous to MBON γ5β′2a terminals. For max-stack: grey, postsynaptic *trans-*Tango signal. **(C)** Heatmap displaying the overlap in segmented MBON postsynaptic signal by brain region. Postsynaptic signal for each MBON was normalized within each brain to capture respective expression levels. SS01308 (MBON γ5β′2a), MB011B (MBONs γ5β′2a, β′2mp, β′2mp_bi), and MB110C (MBONs γ3, β′1) had the strongest postsynaptic signal (max signal 20-24). MB057B (MBON β′1), MB298B (MBON γ4>γ1γ2), MB50B (MBONs α′1, α2sc), SS01143 (MBON β′2mp), and MB433B (MBONs β1>α, γ4>γ1γ2) max signal ranged from 6-11. MB399B (MBON β2β′2a), MB002B (MBONs γ5β′2a, β′2mp), MB077C (MBON γ2α′1), MB018B (MBON α′2), MB027B (MBON α′2), SS01194 (MBON α2sc) max signal ranged from 1-4. For postsynaptic signal normalized across MBON driver lines see Figure 2 Supplementary 1. **(D)** Schematic of fly brain highlighting the most anterior brain regions included in mask analysis starting at AL and ending with SLP. **(E)** Schematic of fly brain highlighting the most posterior brain regions included in mask analysis starting at NO and ending with PB. Abbreviations: AL: antennal lobe, AMMC: antennal mechanosensory and motor center, ATL: antler, AVLP: anterior ventrolateral protocerebrum, CRE: crepine, EB: ellipsoid body, EPA: epaulette, FSB: fan-shaped body, FLA: flange, GA: shoulder of lateral accessory lobe, GOR: gorget of ventral complex, IB: interior bridge, ICL: inferior clamp, IPS: inferior posterior slope, IVLP: inferior ventrolateral protocerebrum, LAL: lateral accessory lobe, LB: bulb of lateral complex, LH: lateral horn, MB: mushroom body, NO: noduli, OTU: optic tubercle, PB: protocerebral bridge, PLP: posteriorlateral protocerebrum, PRW: prow, PVLP: posterior ventrolateral protocerebrum, SAD: saddle, SCL: superior clamp, SEG: subesophageal ganglion, SIP: superior intermediate protocerebrum, SLP: superior lateral protocerebrum, SMP: superior medial protocerebrum, SPS: superior posterior plate, VES: vest of ventral complex, WED: wedge. Scale bar=50μm.

### DANs are postsynaptic to MBONs

Of the DANs innervating the MB, 90% have dendritic arborizations that are localized to 4 of the 5 proposed MBON convergent regions, including CRE, SMP, SIP, and SLP (Aso et al., 2014a). Subsets of MBON axons overlapping with DAN dendritic arborizations provide opportunities for MBONs to modulate DAN activity as well as indirectly modulate their own activity or that of other MBONs. Thus, we selected a subset of MBONs that were reported to co-localize with protocerebral anterior medial (PAM) DANs and co-stained with antibodies against tyrosine hydroxylase (TH) to identify overlap with *trans*-Tango signal (Aso et al., 2014a). As expected, some of the neurons postsynaptic to MBONs were TH positive; however, due to the complexity of *trans-*Tango labeled neurons, we were unable to identify the DANs postsynaptic to a particular MBON unequivocally. Most overlap between TH and *trans*-Tango signals was observed with γ5β′2a (MB011B; Figure 3A) and β′2mp (MB002B; Figure 3B) MBONs. These MBONs were predicted to co-localize with PAM DANs β′2p, β′2m and PAM DANs γ5 and β′2a, respectively (Aso et al., 2014a). Similarly, the cholinergic γ2α′1 MBON (MB077B) was predicted to overlap with PAM γ4<γ1γ2, and indeed, MB077B brains averaged 5 cells with co-expression of TH and *trans-*Tango signals per hemibrain (Figure 3B). Likewise, the γ3, γ3β′1 MBON was predicted to overlap with PAM γ3 and β′1m, and MB083C had an average of 10 cells with co-expression of TH and *trans-*Tango signals (Figure 3C). Despite predictions that γ4>γ1γ2 MBON (MB298B) would co-localize with PAM γ4>γ1γ2, we found minimal co-expression of TH and *trans-*Tango signals. This is likely a false negative due to the strength of the driver as annotations of the EM data has revealed postsynaptic connections with PAM γ4>γ1γ2 (Clements et al., 2020) (Figure 3A). It is possible that the number of co-localized TH+ cells in our analysis here is an underestimation since some of the brains had fewer than expected TH+ neurons (Figure 3 Supplementary 2).

**Figure 3.**
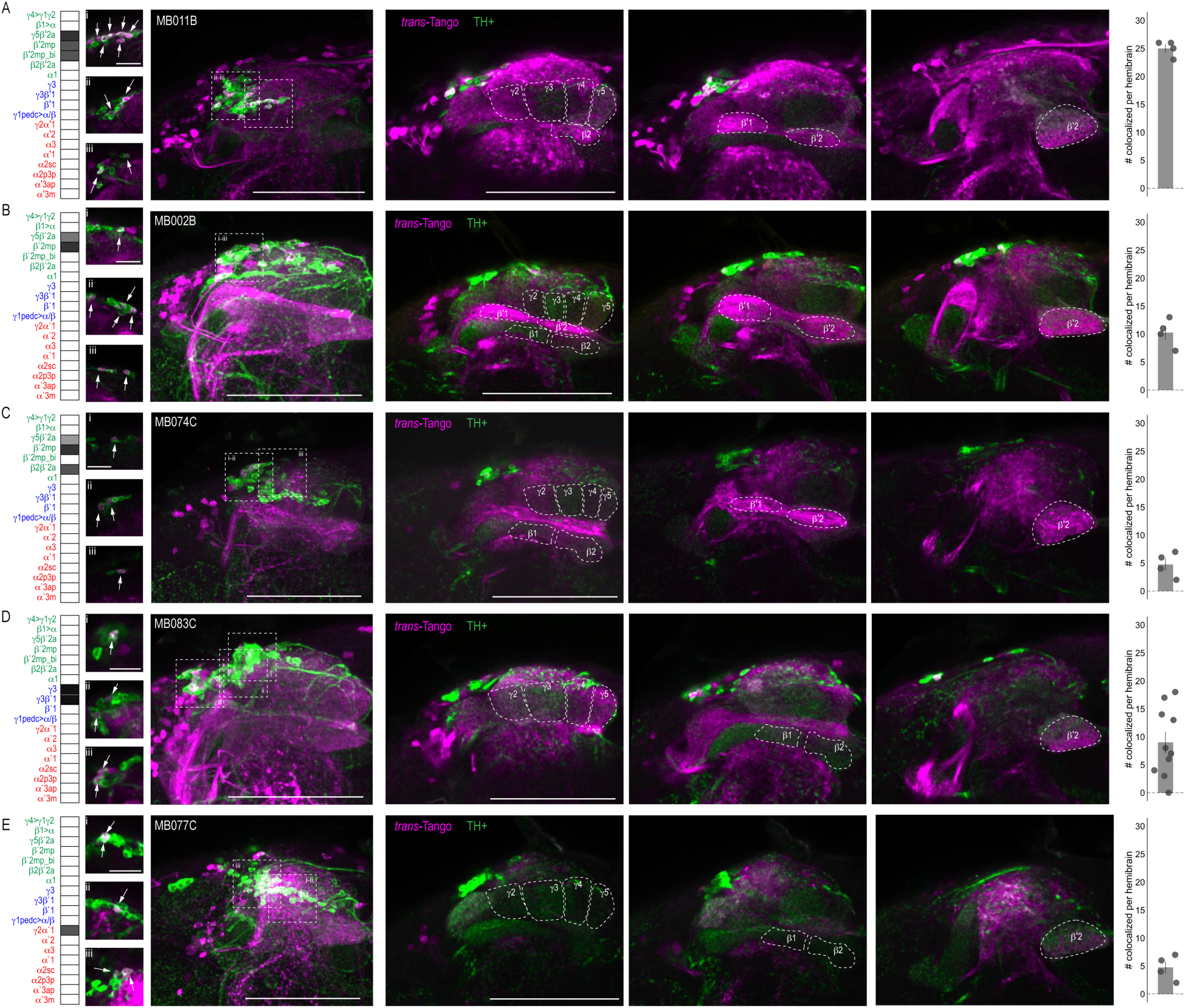
DANs postsynaptic to MBONs. Exemplar max-stacks of MBON lines in which TH+ cells overlapped with postsynaptic signal of glutamatergic **(A)** MBON γ5β′2a, β′2mp, β′2mp_bilateral (MB011B), **(B)** MBON γ5β′2a, β′2mp (MB002B), **(C)** MBON γ5β′2a, β′2mp, β2β ′2a (MB074C), **(D)** GABAergic MBONs γ3, γ3β′1 (MB083C) and **(E)** cholinergic MBONs γ2α′1 (MB077C). Overlapping TH+ and *trans-*Tango cell bodies are highlighted in insets, scale bar =10μm. Max stacks of MB are included, scale bar=50μm. Bar graphs indicate the average number of co-localized cells per hemisphere (mean +/-standard error). Green, TH-positive cells; magenta, postsynaptic *trans*-Tango signal. MBON maps are organized by neurotransmitter type: green=glutamatergic, blue=GABAergic, red=cholinergic.

### Convergent MBONs

Whole brain overlap analysis identified the MB itself as a site of rich convergence for most MBON lines (Figure 2). MBON-postsynaptic signals in MB was not surprising given that many MBONs provide feedforward connections between MB compartments (Aso et al., 2014a). For instance, MBON γ4>γ1γ2 has dendritic arbors in γ4 and axonal projections in γ1γ2, MBON γ1pedc>α/β have dendritic arbors in γ1 and axonal projections in α/β lobes, and MBON β1>α has dendritic arbors in β1 and axon projections to the entire alpha lobe. However, further analysis revealed that in addition to providing connections between MB compartments, MBONs converge directly on other MBONs presumably through axo-axonal connections. Two different MBONs are frequently targeted: MBON β′2mp (Figure 4A) and MBON γ3β′1 (Figure 4B). Interestingly, MBON β′2mp receives convergent glutamatergic, GABAergic, and cholinergic input from MBON γ5β′2a (MB011B and MB210B), MBON γ3β′1 (MB110C and MB83C), MBON α′2 (MB018B and MB082C), and MBON γ2α′1 (MB077B and MB051C) (Figure 4A, Figure 4 Supplementary 1). MBON γ3β′1 receives convergent input from glutamatergic MBON β′2mp as revealed with split-GAL4 lines MB002B (Figure 4B) and MB074C (Figure 4 Supplementary 1) and glutamatergic MBON γ4>γ1γ2 (Figure 4B). We hypothesize that similar to MBONs that project to other regions of the MB, MBON-γ3β′1 and MBON-β′2mp create opportunities for multilevel feedforward networks to update information to drive behavioral response (Figure 4C).

**Figure 4:**
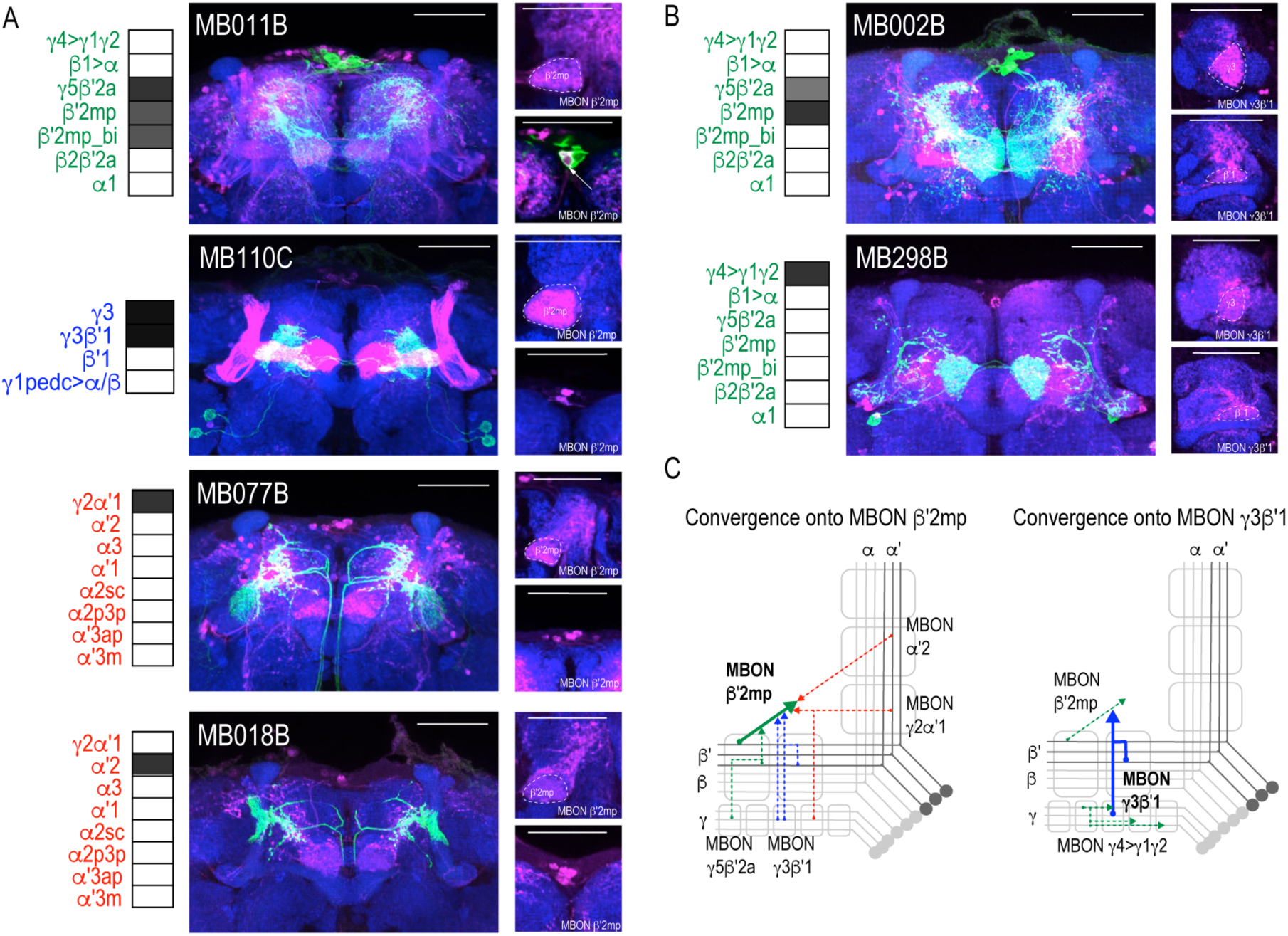
Subsets of MBONs converge on other MBONs. **(A)** MBON β′2mp receives convergent input from glutamatergic MBON γ5β′2a (MB011B), GABAergic MBONs γ3, γ3β′1 (MB110C) and cholinergic MBON γ2α′1 (MB077B) and MBON α′2 (MB018B**). (B)** MBON γ3β′1 receives convergent input from glutamatergic MBON β′2mp (MB002B) and MBON γ4>γ1γ2 (MB298B). β′2mp, γ3 and β′1 are outlined in representative stacks. **(C)** Schematics summarizing identified convergent MBONs (β′2mp and γ3β′1) and their respective convergent input. Solid lines represent the convergent MBON and dotted lines represent convergent input. For max-stacks: green, presynaptic MBONs, magenta, postsynaptic *trans*-Tango signal, blue, *brp-SNAP* neuropil, scale bar=50μm.

### Convergence outside the MB

Another site of convergence of the MBON network was the FSB (Figure 5). MBON postsynaptic connections display a laminar organization primarily across the dorsal region of the FSB. Nearly all of the glutamatergic and GABAergic MBONs converge onto FSB layers 4 and 5, and to a lesser extent, layer 6 (Figure 5A-C). MBON α1 is the only type of MBON that had broad *trans*-Tango signals in the FSB (Figure 5B). Cholinergic MBONs also had *trans*-Tango signals in the dorsal FSB but with more variability across MBON lines and within each line. For instance, *trans*-Tango with MBON γ2α′1 consistently visualized projections to FSB layers 4 and 5 in all of the brains analyzed, whereas more variability was observed in FSB innervation pattern across MBON α′2 brains (Figure 5 Supplementary 1). MBON α′1 and α2sc both project exclusively to FSB layer 6 (Figure 5C). Together, FSB layers 4 and 5 receive convergent input from combinations of glutamatergic, GABAergic and cholinergic MBONs (Figure 5D).

**Figure 5.**
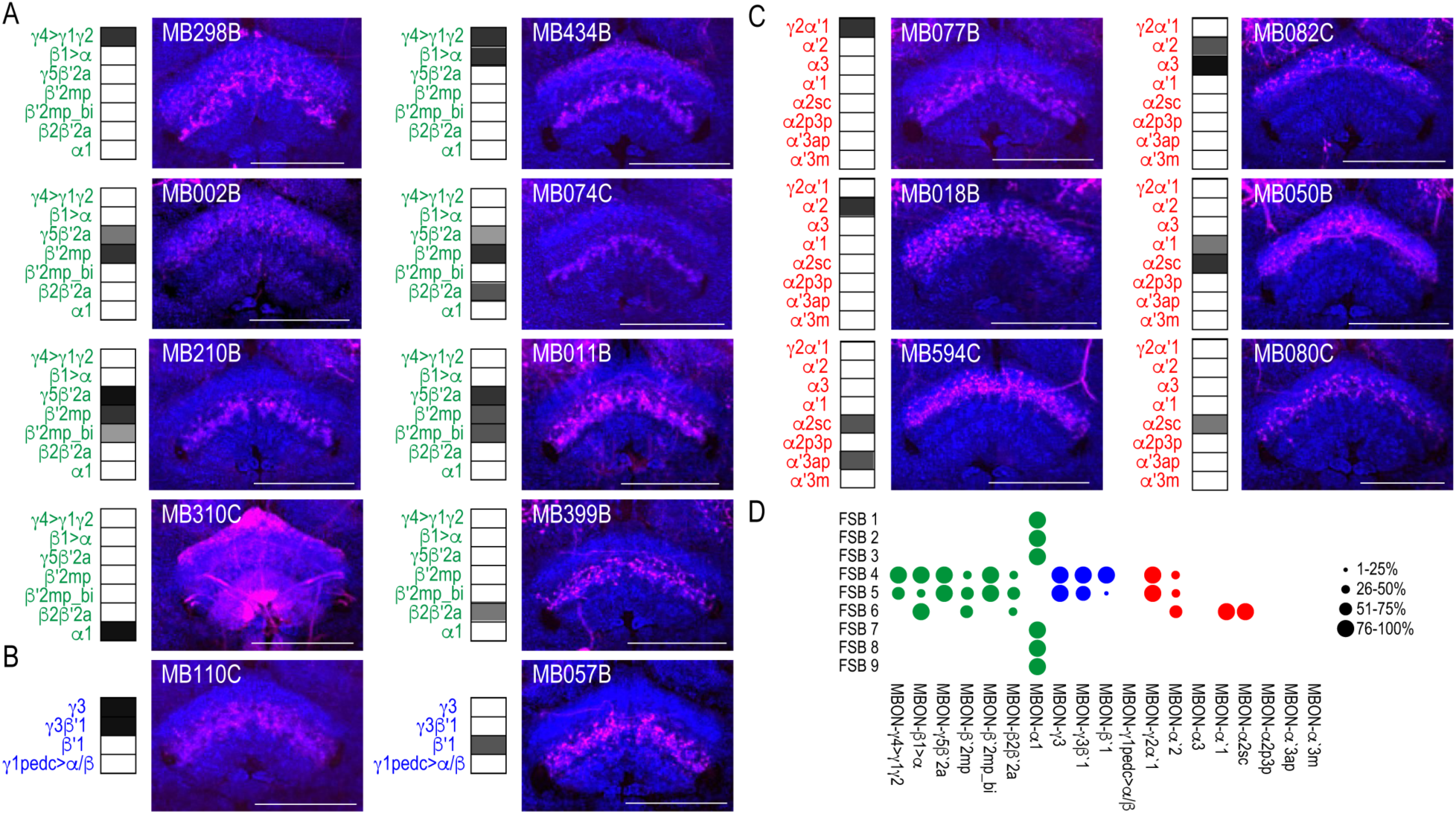
MBONs converge on different layers of the FSB. Exemplar max-stacks of glutamatergic **(A)**, GABAergic **(B)**, and cholinergic **(C)** MBONs whose postsynaptic neurons innervate the FSB. Max-stacks are approximately 50μm thick. Slices were selected based on the relative position of the FSB. For FSB stacks: magenta, postsynaptic *trans*-Tango signal, blue, *brp-SNAP* neuropil. Map of MBONs accompany each exemplar with the relative expression pattern (greyscale, 1-5) accordingly to FlyLight. For each map green=glutamatergic, blue=GABAergic, red=cholinergic. Scale bar=50 μm. **(D)** Map summarizing the percentage of *trans-*Tango positive signal in each FSB layer across brains for each MBON.

Both visual and computational analyses identified the CRE, SMP, SIP, and SLP, as well as the MB and FSB as obvious postsynaptic targets of the MBON network. Visual inspection also identified the LAL as postsynaptic to multiple MBON lines. Its identification was less obvious in computational analysis largely because the neurites innervating the LAL were not as extensive as the LAL itself and were often difficult to segment. Although not extensive, LAL innervation was consistent across glutamatergic, GABAergic, and cholinergic MBONs (Figure 6). Specifically, glutamatergic γ5β′2a, β′2mp, and β′2mp_bilateral had postsynaptic neurites within the LAL in all of the brains analyzed (Figure 6A). Similarly, GABAergic MBON γ3, γ3β′1, and β′1 (Figure 6B) and cholinergic MBON γ2α′1 (Figure 6C) consistently had postsynaptic neurites within the LAL. Thus, like the FSB, the LAL receives convergent input from combinations of glutamatergic, GABAergic and cholinergic MBONs (Figure 6D).

**Figure 6.**
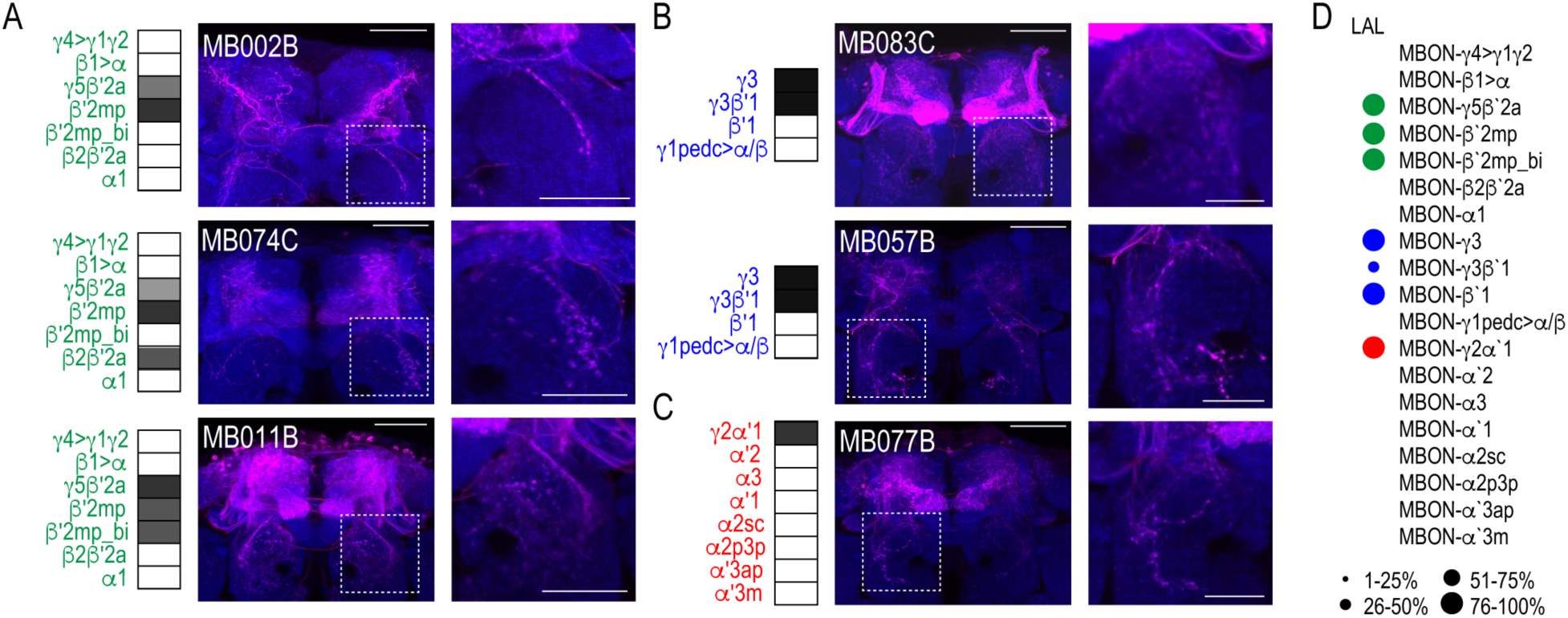
MBONs converge onto LAL neurons. Exemplar max-stacks of glutamatergic **(A)**, GABAergic **(B)**, and cholinergic **(C)** MBONs innervating the LAL. Max-stacks are approximately 50μm thick. Slices were selected based on the relative position of the LAL. Magenta, postsynaptic *trans*-Tango signal, blue, *brp-SNAP* neuropil. Map of MBONs accompany each exemplar with the relative expression pattern (greyscale, 1-5) accordingly to FlyLight. For each map green=glutamatergic, blue=GABAergic, red=cholinergic. Scale bar=50μm. Scale bar for insets = 10μm **(D)** Map summarizing the percentage of *trans-*Tango positive signal in LAL across brains for each MBON.

Thus far, we have identified two postsynaptic targets of the MBON network that reside outside of the MB: the FSB and LAL. However, the identities of the postsynaptic neurons within FSB and LAL as well as their functions remain unknown. Our strategy for identifying FSB and LAL neurons and interrogating their functional connectivity with MBONs was to selectively label neurons in FSB and LAL using specific drivers and to examine whether they are co-localized with postsynaptic signal when we initiate *trans*-Tango from MBONs. To achieve this, we identified candidate FSB and LAL LexA lines by performing a mask search of the LexA lines that have overlapping expression within the convergent region and brought them together with MBON lines: MB051C and MB077C were used to target MBON γ2α′1, MB083C and MB110C were used to target γ3β′1, and MB074C was used to target MBON β′2mp. We identified three candidate LexA lines: one to target FSB layer 4 neurons -R47H09 (Jenett et al., 2012; Pfeiffer et al., 2013.10.9; Pfeiffer et al., 2010), and two to target LAL neurons -VT055139 and VT018476 (Tirian and Dickson, 2017). Finally, we generated *trans-*Tango reporter flies where the UAS-myrGFP was replaced with UAS-CD2, and LexAOp-mCD8::GFP was included in order to visualize the starter MBONs, the postsynaptic *trans*-Tango signal, and the LexA lines simultaneously.

We successfully combined the majority of the targeted MBON split-Gal4 lines with FSB and LAL LexA lines (we were unable to combine MB074C with LexA line 47H09). Interestingly, for the cholinergic MBON γ2α′1, we identified at least two postsynaptic FSB neurons (labeled in the 47H09 LexA line; Figure 7A) and at least five postsynaptic LAL neurons (labeled in the VT055139 LexA line; Figure 7B). We next sought to interrogate functional connectivity between MBON γ2α′1 and 47H09 FSB neurons and VT055139 LAL neurons by combining optogenetic stimulation of MBON γ2α′1 and functional calcium imaging in FSB and LAL. To achieve this, we combined reporter flies (UAS-Chrimson and LexAop-GCaMP6s) with MB077C. Stimulation of cholinergic MB077C with 400-500ms of red light (627 nm) resulted in an increase in calcium signal in the FSB and LAL (Figure 7C). Similar activation of other cholinergic MBONs, which do not innervate the LAL or layer 4 of the FSB (MB080C), did not result in signal (Figure 7 Supplementary 1), supporting the specificity of this interaction and suggesting that the MBON γ2α′1 is both anatomically and functionally connected to the FSB and LAL. Strikingly, GABAergic MBON γ3β′1 also had at least one identified postsynaptic FSB neuron that was included in the expression of FSB 47H09 LexA line (Figure 7D) and at least two identified postsynaptic LAL neurons that were included in the expression of LAL VT055139 LexA line (Figure 7E). Thus, the genetically identified subsets of LAL and FSB neurons receive convergent input from GABAergic and cholinergic MBONs (Figure 7F). We hypothesize that the convergence of excitatory and inhibitory input onto both the LAL and FSB is critical for guiding behavior.

**Figure 7.**
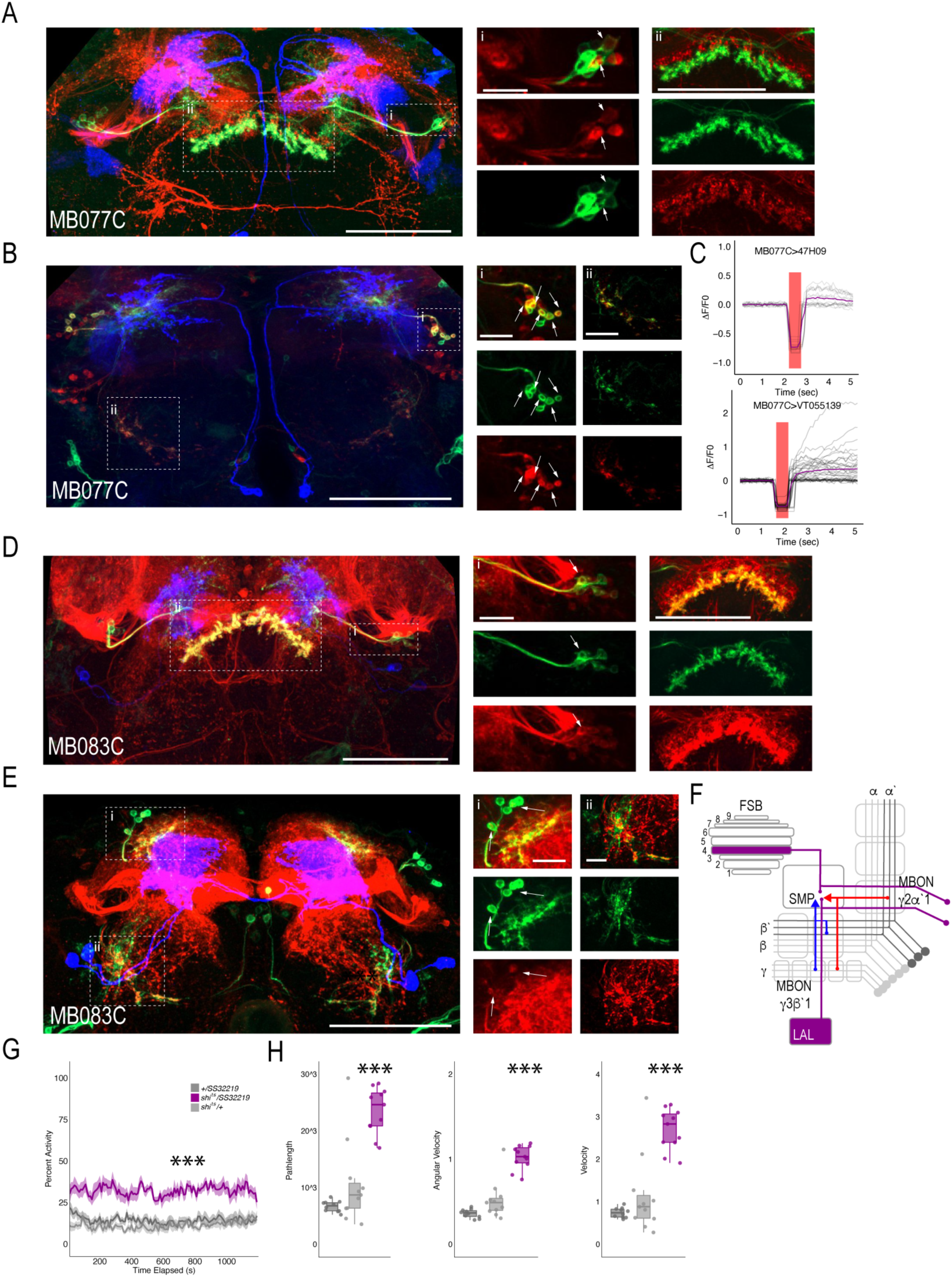
MBONs γ3β′1 and γ2α′1 converge onto the same subset of LAL and FSB neurons. Exemplar max-stacks of cholinergic MBON γ2α′1 postsynaptic connections and identified overlap with respective **(A)** FSB (47H09) and **(B)** LAL (VT015539). **(C)** Confirmation of functional connection with optogenetic activation of MB077C and calcium imaging of FSB neurons in SMP and FSB (47H09), and calcium imaging of LAL neurons in SMP (VT015539). The red bar indicates when the LED was on and the shutter was closed to protect the PMTs during LED stimulation. Exemplar max-stacks of GABAergic MBON γ3β′1 postsynaptic connections and identified overlap with respective **(D)** FSB (47H09) and **(E)** LAL (VT015539). Max-stacks are approximately 50μm thick. Slices were selected based on the relative position of the LAL and FSB. In A, B, D and E, red, postsynaptic *trans*-Tango signal; blue, CD2 marker of split-GAL4 line; green, LexA FSB or LAL. Scale bar=50μm. **(F)** Schematic highlighting convergence of MBONs γ3β′1 and γ2α′1 onto the same genetically identified subsets of LAL and FSB neurons. **(G)** *shibire*^ts^ inactivation of LAL using split-GAL4 SS32219 resulted in significant increases in group activity (F(2,21)=39.28 p<0.0001). Group activity counts were binned over 10s periods, averaged across biological replicates of 10 flies each (n=8) and plotted against time. Lines depict mean +/-standard error. **(H)** One video was selected at random of each genotype and processed using FlyTracker to calculate the average pathlength (F(2,29)=33.39, p<0.0001), angular velocity (F(2,29)=51.87, p<0.0001) and velocity (F(2,29)=30.97, p<0.0001) of individual flies. Box plots with overlaid raw data were generated using RStudio. Each dot is a single fly. One-way ANOVA with Tukey Posthoc was used to compare mean and variance. ***p<.0001.

Finally, to determine the role of LAL neurons in the context of guiding behavior of flies in groups, we performed analyses of group activity using thermogenetic inactivation of identified split-GAL4 LAL neurons (Scaplen et al., 2019). Individual flies were tracked offline using Flytracker to obtain activity based features (Eyjolfsdottir et al., 2014). Inactivation of SS32219-GAL4 positive LAL neurons resulted in significant increases in group activity (Figure 7G, (F(2,21)=39.28 p<0.0001), pathlength (F(2,29)=33.39, p<0.0001), angular velocity (F(2,29)=51.87, p<0.0001) and velocity (F(2,29)=30.97, p<0.0001) of individual flies (Figure 7H). Behavioral results were replicated with a separate LAL split-GAL4 line (SS32230-GAL4, Figure 7 Supplementary 2), suggesting that LAL neurons downstream of MBONs modulate locomotor activity of flies in a group. Group activity at permissive temperatures was not different from controls (Figure 7 Supplementary 3).

## Discussion

The MB is a high-level integration center in the *Drosophila* brain with an established role in learning and memory. The iterative nature of converging and diverging MB neural circuits provides an excellent example of the anatomical framework necessary for complex information processing. For instance, on a rapid timescale, interactions between MB compartments could generate different output patterns to drive behavior, whereas on a slower timescale, interactions between MB compartments could reevaluate memories of a context (Aso and Rubin, 2016; Felsenberg et al., 2017; Felsenberg et al., 2018).

We sought to map divergence and convergence of the neural circuits beyond the MB using the genetic anterograde transsynaptic technique, *trans-*Tango. Our study complements the ongoing efforts of EM studies (Li et al., 2020; Xu et al., 2020) for the whole brain of a single female fruit fly, as we report connectivity of MBONs across multiple subjects in both males and females, and highlight the variability in connectivity that potentially exists across animals. Although the complete EM dataset of an adult fly brain has been an invaluable resource that significantly accelerated the mapping of the neural circuits underlying innate and learned behaviors (Adden et al., 2020; Li et al., 2020; Xu et al., 2020; Zheng et al., 2018), the massive undertaking of acquiring a full EM dataset renders it impractical to perform for multiple individuals. The examination of the circuit connectivity of several individuals, easily afforded by the use of *trans*-Tango, expands the value of the EM reconstruction data by adding nuanced differences between individuals. Further, *trans*-Tango can be readily adapted to functional studies in which the activity of the postsynaptic neurons is altered by expressing optogenetic/thermogenetic effectors or the activity of the postsynaptic neurons is monitored by expressing genetically encoded sensors. Our tracing studies reported here serve as the foundation for these future experiments.

Our studies reveal that the MB circuits are highly interconnected with multiple regions of converging projections both within and downstream of the MB. Our experiments also reveal diverging projections in the downstream postsynaptic targets. We anatomically and functionally describe a multilayer circuit that includes GABAergic and cholinergic MBONs that converge on the same subset of FSB and LAL neurons. This circuit architecture provides an opportunity to rapidly update online processing of sensory information before executing behavior.

### Divergence across the brain

Successive levels of convergence and divergence across the brain permit functional flexibility (Jeanne and Wilson, 2015; Man et al., 2013).(Tye, 2018). Our data revealed varying levels of divergence of postsynaptic connections of MBONs across the brain. Every one of the analyzed MBONs had postsynaptic partners projecting to multiple brain regions (Figure 2C). Further, nearly the entire superior protocerebrum as well as portions of the inferior protocerebrum received input from at least one MBON, providing opportunities for comprehensive integration of signals from the MBON network.

### Convergence within the MB

Multiple feedforward and feedback circuits exist within the MB (Aso et al., 2014a; Eichler et al., 2017; Eschbach et al., 2020; Otto et al., 2020; Takemura et al., 2017; Zheng et al., 2018). Our data revealed at least two MBONs that receive convergent input from multiple MBONs and are also reciprocally connected. MBON β′2mp receives convergent input from glutamatergic MBON γ5β′2a, cholinergic MBONs γ2α′1 and α′2 as well as from the GABAergic MBON γ3β′1, which also receives convergent input from other MBONs (Figure 3). The convergent MBON input to MBON β′2mp is especially interesting as these cholinergic, GABAergic, and glutamatergic MBONs drive opposing behaviors (Aso et al., 2014b). For instance, activation of the cholinergic MBONs γ2α′1 or the GABAergic MBON γ3β′1 results in naïve preference, whereas activation of the glutamatergic MBON γ5β′2a results in robust naïve avoidance (Aso et al., 2014b; Lewis et al., 2015). Similarly, MBON γ2α′1 mediates aversive associations (Berry et al., 2018; Yamazaki et al., 2018), whereas MBON γ5β′2a mediates appetitive associations and extinction of aversive memories (Owald et al., 2015) (Yamazaki et al., 2018) (Felsenberg et al., 2018). Considering that MBON β′2mp receives convergent input from these parallel and opposing pathways, it likely serves as a decision hub by integrating activity to modulate cue-induced approach and avoidance behavior. Further, since the roles of these converging MBONs in naïve and learned behaviors are state dependent (Grunwald Kadow, 2019; Tsao et al., 2018) (Lewis, 2015 #78}, we hypothesize that MBON-γ3β′1 and MBON-β′2mp provide opportunities for multilevel feedforward networks to update information processing.

Some of the feedback connections in the MB originally hypothesized to exist were between MBONs and DANs innervating the MB (Aso et al., 2014b; Ichinose et al., 2015). Our analysis revealed neurons postsynaptic to MBONs that are TH positive. Recent studies that use a combination of EM annotation and calcium imaging to identify specific MBON-DAN connections suggest extensive recurrent connectivity between MBONs and DANs, validating our findings (Felsenberg et al., 2018; Li et al., 2020; Otto et al., 2020). For example, previous studies using both GRASP and EM annotation revealed that MBON α1 and PAM α1 are synaptically connected (Ichinose et al., 2015) (Li et al., 2020). We similarly identified a few PAM neurons within the MBONα1 postsynaptic signal. A recent study showed that the 20 DANs that innervate the γ5 MB compartment are clustered into 5 different subtypes that innervate distinct anatomical regions within the γ5 compartment (Otto et al (2020). According to this study, only one of the PAM γ5 DANs receives direct recurrent feedback from γ5β′2a MBONs (Otto et al., 2020). Based on these recent anatomical characterizations, we believe that the TH+ neurons within the postsynaptic signal of γ5β′2a are the γ5 DANs.

### Convergence within the FSB

The FSB is the largest substructure of the central complex and it serves as a sensory-motor integration center (Pfeiffer and Homberg, 2014; Wolff et al., 2015). The FSB comprises 9 horizontal layers (Wolff et al., 2015) that are innervated by large-field neurons (Hanesch et al., 1989). Our data suggest that the large-field neurons of the dorsal FSB are postsynaptic to the majority of MBONs. Although there exists some variation across brains, glutamatergic and GABAergic MBONs predominately project to FSB layers 4 and 5, whereas cholinergic MBONs mainly project to FSB layer 6. Connections between MBONs and FSB were consistent across different split-GAL4 lines that have overlapping expression patterns. Similar extensive direct connectivity between these MBONs and the dorsal FSB, especially layers 4 and 5, were found in the recently annotated EM hemibrain dataset (Li et al., 2020).

How are FSB layers 4/5 and 6 functionally distinct? The dorsal FSB has a well-established role in modulating sleep and arousal (Berry et al., 2015; Donlea et al., 2011; Ueno et al., 2012), locomotor control (Strauss, 2002), courtship (Sakai and Kitamoto, 2006) and visual memory (Li et al., 2009; Liu et al., 2006; Wang et al., 2008). FSB layer 5 has been specifically implicated in processing elevation in a *foraging-* and *rutabaga-*dependent manner (Li et al., 2009). More recent studies have implicated the dorsal FSB in processing nociceptive information (Hu et al., 2018). FSB layer 6 plays a specific role in avoidance of a conditioned odor, whereas layers 4 and 5 respond to aversive stimuli and are responsible for innate, but not conditioned, avoidance (Hu et al., 2018). Moreover, recent connectome data suggest that differences exist in the postsynaptic connections of layers 4/5 and 6 as well. Overall there is high degree of interconnectivity within the FSB (Clements et al., 2020). The predominate output of FSB layer 6 neurons are other FSB neurons. In fact, many FSB layer 6 neurons project exclusively to other FSB neurons (Clements et al., 2020). In contrast, FSB layer 4 neurons send direct projections to CRE, SMP, and LAL in addition to projecting to other FSB neurons. The connections with the LAL position the FSB layer 4 to directly influence downstream motor output signals prior to executing behavior. Recent EM analysis also suggests that some FSB layer 6 neurons synapse back onto PAM DAN neurons (Li et al., 2020). This connectivity is in line with the associative role in conditioned nociception avoidance described for FSB layer 6 (Hu et al., 2018).

Interestingly, we found the pattern of FSB postsynaptic targets of the MBONα1 is dissimilar to other glutamatergic MBONs. FSB layers 4/5 and 6 are not present in the MBONα1 postsynaptic signal. Instead, MBONα1 project to neurons that innervate the ventral and most dorsal aspect of the FSB. The ventral FSB is implicated in innate avoidance of electric shock (Hu et al., 2018), and more recent data suggest that its activity is tuned to airflow cues for orientation during flight (Currier et al., 2020). Artificial activation of MBONα1 does not result in significant avoidance behavior (Aso et al., 2014b). However, it has been implicated in the acquisition, consolidation, and expression of 24hr long-term sucrose memory (Ichinose et al., 2015). It is possible that MBONα1 provides appetitive valence signals to the ventral FSB to guide goal directed flight. Functionally validating the role of MBONα1 and its relationship with its putative downstream neurons is key to appreciating how learning signals can drive behavioral decisions.

More research is necessary to further understand the functional role of different FSB layers and how information is integrated across these layers. Based on the anatomical data, it is clear that although the MB and FSB can function in parallel during memory formation, they act as parts of a dynamic system to integrate information and adjust behavioral responses.

### Convergence within the LAL

The LAL is an important premotor waystation for information traveling from the central complex to descending neurons innervating thoracic motor centers across insects (Chiang et al., 2011; Franconville et al., 2018; Hanesch et al., 1989; Namiki and Kanzaki, 2016; Wolff and Strausfeld, 2015). Accordingly, the LAL has been implicated in orientation to pheromones (Kanzaki et al., 1991a, b; Mishima and Kanzaki, 1999; Namiki et al., 2014; Namiki et al., 2018), flight (Homberg, 1994), locomotion (Bidaye et al., 2014) and in response to mechanosensory stimuli (Homberg, 1994). More recent work has suggested a functional organization whereby the neurons in the upper division of the LAL receive convergent input from the protocerebrum and neurons in the lower division generate locomotor command (Namiki et al., 2014; Rayshubskiy et al., 2020).

Our data show that the MB network converges with the protocerebrum input, thereby providing an opportunity for MBONs to indirectly influence descending motor outputs. We also demonstrate that two MBONs (γ3β′1 and γ2α′1) synapse on the same subset of LAL and FSB cells, revealing a convergent circuit that connects both structures. Further, in support of our anatomical observations, optogenetic activation of MBON γ2α′1 resulted in activation of both LAL and FSB layer 4 neurons. Given that MBON γ3β′1 is GABAergic, we did not perform the equivalent experiment for this neuron. Thus, the functional consequences of these inhibitory connections remain to be seen. Interestingly, despite the fact that MBON γ3β′1 and γ2α′1 express different neurotransmitters and innervate different MB compartments, their manipulation has similar behavioral phenotypes: both promote sleep (Aso et al., 2014b; Sitaraman et al., 2015a; Sitaraman et al., 2015b), and artificial activation of either results in naïve preference (Aso et al., 2014b). Further, activation of both MBON γ3β′1 and γ2α′1 together has an additive effect, which results in a significant increase in preference (Aso et al., 2014b).

The behavioral significance of the projections of MBON γ3β′1 and γ2α′1 to both the FSB and LAL remains to be determined. We found that inactivation of the putative downstream LAL neurons significantly increased overall activity of behaving flies in a social context and locomotor assay. Since some LAL neurons make direct connections to descending neurons that control walking, this framework can provide a complete circuit through which sensory signals are integrated with punishment or reward to direct the motion of the animal (Rayshubskiy et al., 2020). That said, as the LAL innervation is complex (Franconville et al., 2018), the behavioral implications could be state dependent (Homberg, 1994; Namiki and Kanzaki, 2016).

Understanding how memories are formed, stored, and retrieved necessitates knowledge of the underlying neural circuits. Our characterization of the architecture of the neural circuits connecting the MB with downstream central complex structures lays the anatomical foundation for understanding the function of this circuitry. Our studies may also provide insight into general circuitry principles for how information is processed to form memories and update them in more complex brains.

## Material and Methods

### Fly Strains

All *Drosophila melanogaste*r lines were raised at 18°C on standard cornmeal-agar media with tegosept anti-fungal agent and in humidity-controlled chambers under 14/10 hr light/dark cycles. For a list of fly lines used in the study, see the Key Resource Table.

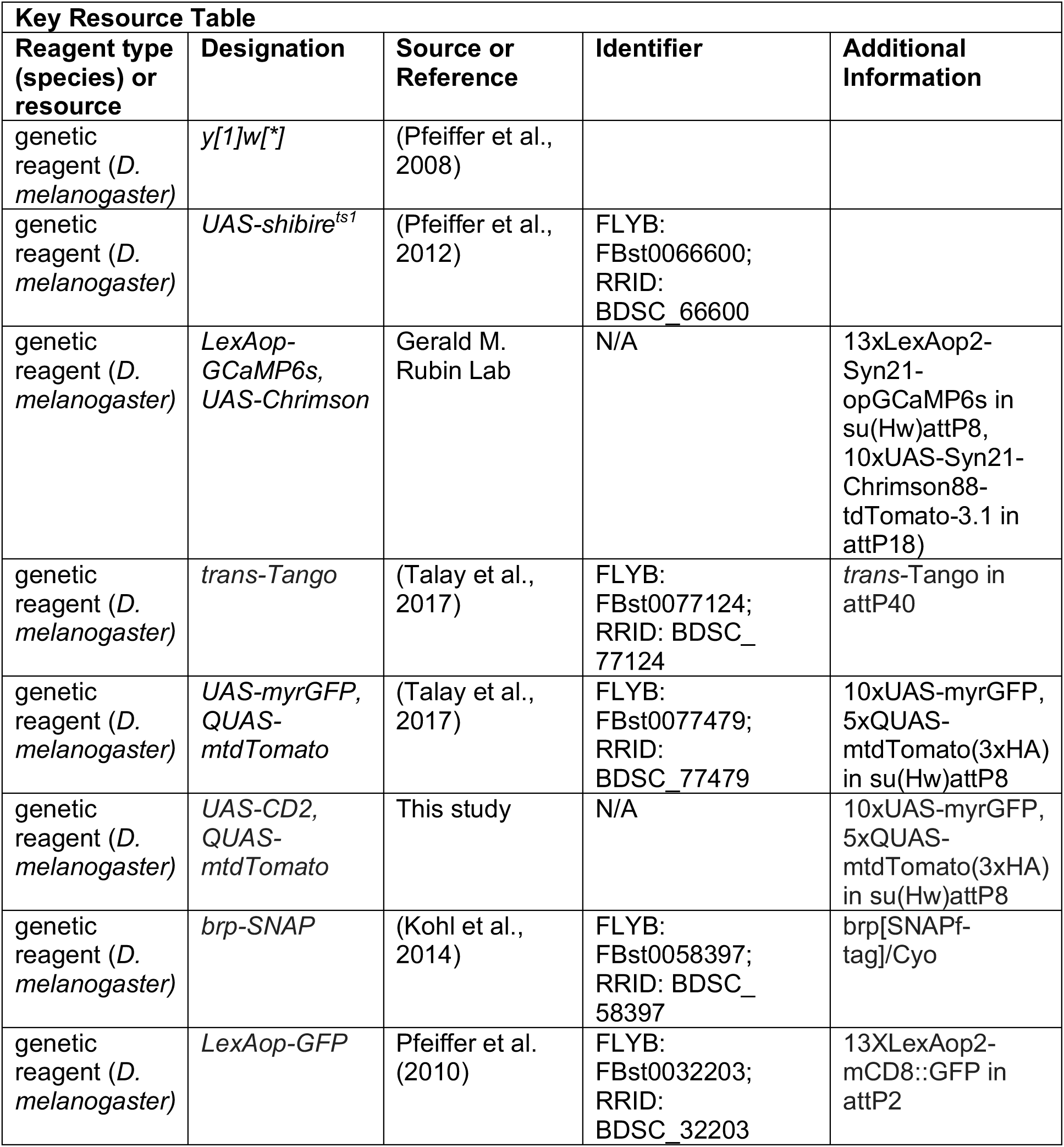
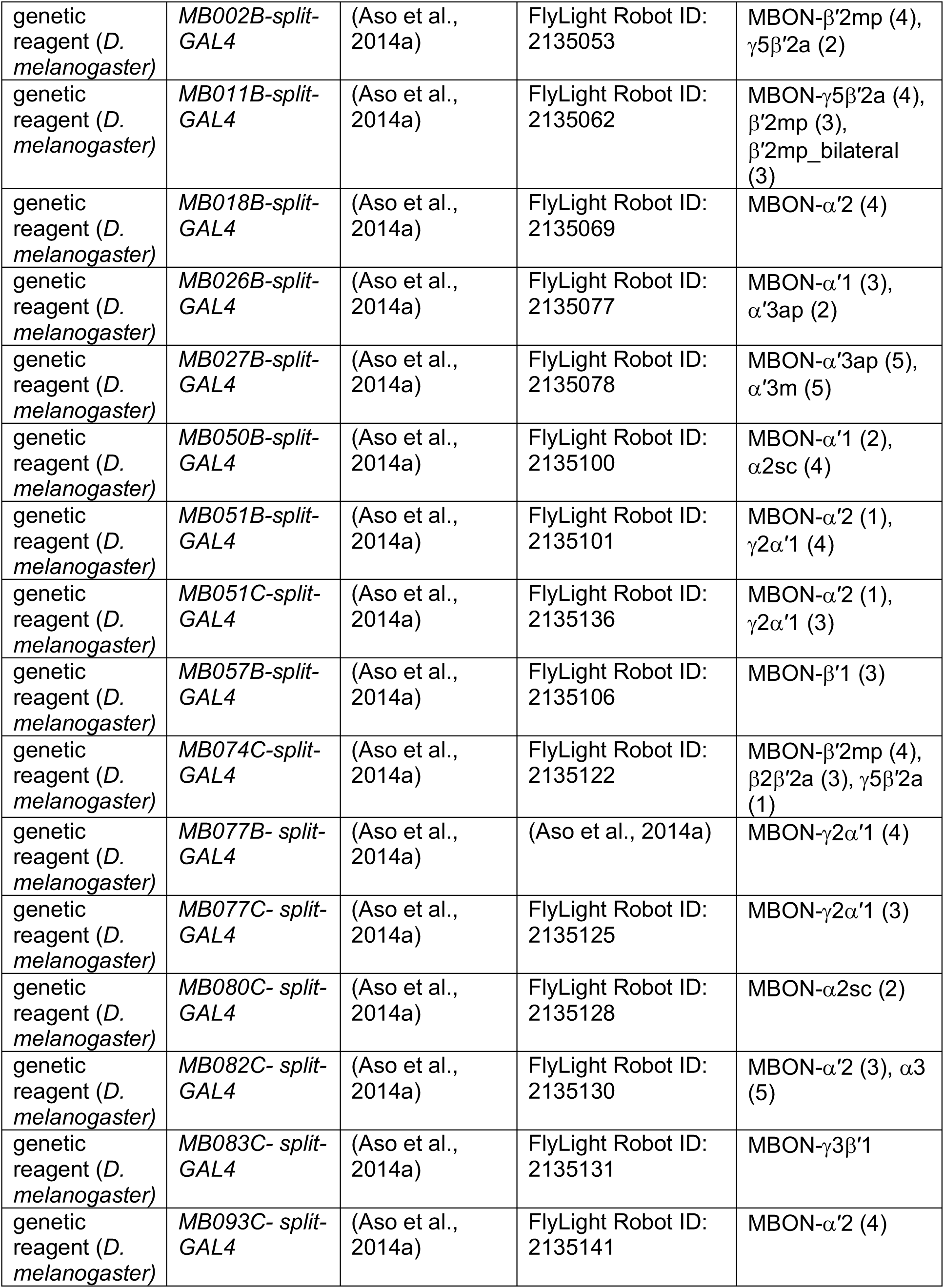
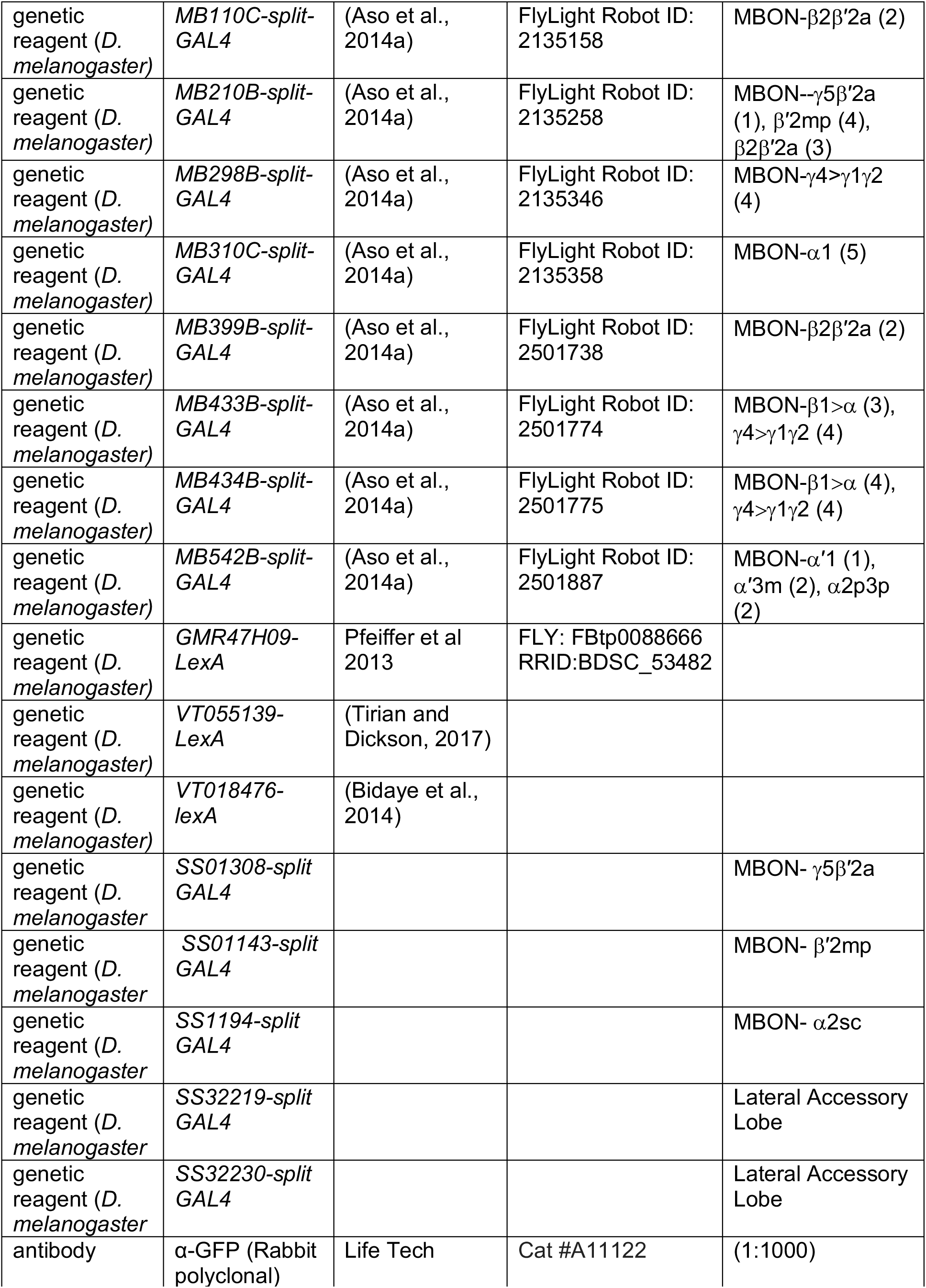
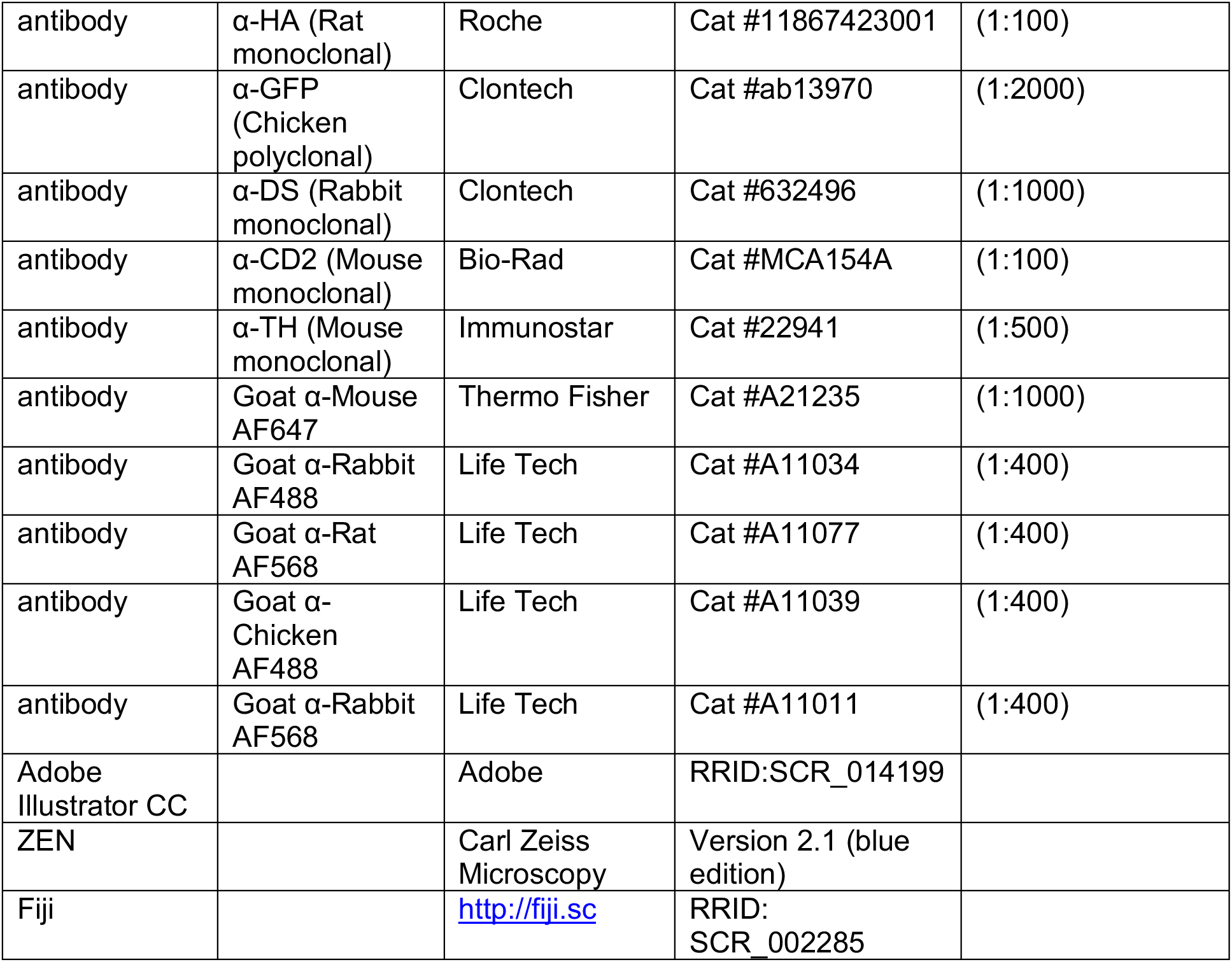

### Generation of transgenic UAS-CD2, QUAS-mtdTomato lines

Gibson Assembly was used to generate the plasmid UAS-CD2_QUAS-mtdTomato(3xHA). The DNA sequence encoding Rattus norvegicus CD2 (NP_036962.1) was codon optimized for Drosophila melanogaster and synthesized by Thermo Fisher Scientific, USA. This sequence was subsequently amplified using primers 5’-atcctttacttcaggcggccgcggctcgagaatcaaaATGCGCTGCAAGTTCCTG-3’ and 5’-agtaaggttccttcacaaagatcctctagaTTAGTTGGGTGGGGGCAG-3’ to obtain the insert fragment. To generate the vector fragment, the *trans*-Tango reporter plasmid (UAS-myrGFP_QUAS-mtdTomato(3xHA)) (Talay et al., 2017) was digested with XhoI and XbaI. Insert and vector fragments were ligated using HiFi DNA Assembly Kit (New England Biolabs, USA) following manufacturer’s instructions. The resultant plasmid was integrated at the su(Hw)attP8 site via PhiC31-mediated recombination and flies were combined with LexAOp-GFP in attP2.

### trans-Tango immunohistochemistry

Flies were dissected at 15-20 days post eclosion using methods adapted from FlyLight Protocols (https://www.janelia.org/project-team/flylight/protocols). Flies were anesthetized with temperature, dewaxed in 70% ethanol, rinsed in Schneider’s Insect Medium (S2) and dissected on a Sylgard pad with cold S2. Within 20 minutes of dissection, collected brains were transferred to 2% paraformaldehyde (PFA) in S2 at room temperature and incubated for 55 minutes. After fixation, brains were rinsed with phosphate buffered saline with 0.5% Triton X-100 (PBT) for 15 minutes at room temperature. Washes were repeated 4 times before storing the brains overnight in 0.5% PBT at 4°C. For chemical tagging in brp-SNAP+ brains, PBT was removed and SNAP substrate diluted in PBT (SNAP-Surface649, NEB S9159S; 1:1000) added. Brains were incubated for 1 hour at room temperature and rinsed with PBT (3 times for 10 minutes). Brains were then blocked in 5% GS (Goat Serum) diluted in PBT for 90 minutes at room temperature. Brains were then incubated in primary antibodies diluted in 5% GS/PBT for 4 hours at room temperate and then at 4°C for two overnights. After primary antibody incubation, brains were washed four times for 10 minutes with 0.5% PBT before incubating in second antibody diluted in 5% GS/PBT at 4°C for two overnights. Samples were then rinsed and washed 4 times for 15 minutes in 0.5% PBT at room temperature and prepared for DPX mounting. Briefly, brains were fixed a second time in 4% PFA in PBS for 4 hours at room temperature and then washed 4 times in PBT for 15 minutes at room temperature. Brains were rinsed for 10 minutes in PBS, placed on PLL-dipped cover glass, and dehydrated in successive baths of ethanol for 10 minutes each. Brains were then soaked 3 times in xylene for 5 minutes each and mounted using DPX.

### Genetic overlap analysis

MBON split-GAL4 “C” lines which have the DNA binding domain (attP2) and activation domain (VK00027) recombined on the 3rd chromosome were crossed to newly generated *trans-*Tango reporter flies where the UAS-myrGFP was replaced with 10xUAS-CD2, and 13xLexAOp-mCD8::GFP was inserted into attP2. This enabled the visualization of the starter MBONs, the postsynaptic *trans*-Tango signal, and the LexA lines simultaneously.

### Microscopy and Image Analysis

Confocal images were obtained using a Zeiss, LSM800 (Brown University) and LSM710 (Janelia Research Campus) with ZEN software (Zeiss, version 2.1) with auto Z brightness correction to generate a homogeneous signal and were formatted using Fiji software (http://fiji.sc). Whole brains were scanned using a 40x objective in four overlapping tiles and then stitched together in the ZEN software.

TH+ cells, and cells with overlapping TH and *trans*-Tango signal were counted by blinded experimenter using the Cell Counter plugin in FIJI (https://imagej.net/Cell_Counter). We counted the total number of TH+ cells that co-localized with *trans*-Tango labeled cells in each hemibrain starting at the most anterior surface of the brain and continued to count TH+ cells until we reached the protocerebral anterior lateral (PAL) cluster which were identified by their cell body size. We did not identify any co-localized cells within or posterior to the PAL cluster.

Images were prepared for publication in FIJI and Adobe Illustrator with no external manipulation aside from cropping to demonstrate higher resolution. All figures were generated using Adobe Illustrator CC.

### Brain registration and tracing postsynaptic connections

Brains were registered as previously described (Aso et al., 2014a). Postsynaptic connections of registered brains were segmented in VVDViewer (https://github.com/takashi310/VVD_Viewer) and saved as .nrrd files. Segmented files of postsynaptic signal for each MBON were multiplied by 34 binary masks of each central brain region in a custom written Matlab program to calculate the distribution of postsynaptic signal across brain regions. Heatmaps were generated in RStudio.

### Calcium imaging protocol and analysis

All functional imaging experiments were performed *ex-vivo* from brains of 1-4-day old male or female brains on an Ultima two-photon laser scanning microscope (Bruker Nanosystems) equipped with galvanometers driving a Chameleon Ultra II Ti-Sapphire laser. Images were acquired with an Olympus 60x, 0.9 numerical aperture objective at 512×512 pixel resolution.

Flies were placed on food containing 400uM all trans-retinal for 18-36 hours prior to dissection. Brains were dissected in saline (108 mM NaCl, 5mM KCL, 2mM CaCl2, 8.2 mM MgCl2, 4mM NAHCO3, 1 mM NaH2PO4, 5mM trehalose, 10mM sucrose, 5mM HEPES, pH 7.5 with osmolarity adjusted to 275 mOsm), briefly (45 s) treated with collagenase (Sigma #C0130) at 2mg/mL in saline, washed, and then pinned with fine tungsten wires in a thin Sylgard sheet (World Precision Instruments) in a 35 mm petri dish (Falcon) filled with saline. MBONs were stimulated with 400ms-500ms of 627 nm LED. For recordings in the LAL (VT018476 and VT055139) ROI were positioned over SMP. For recordings in the FSB (476H09) ROIs were positioned over SMP or FSB.

All image processing was done using FIJI/ImageJ (NIH). Further analysis was performed using custom scripts in ImageJ, Microsoft Excel, and RStudio. Normalized time series of GCaMP fluorescence were aligned to the time point when the opto-stimulus was applied for each replicate.

### Behavioral experiments

Locomotor activity was evaluated in a 37mm diameter circular open field area as described previously (Scaplen et al., 2019). Briefly, for thermogenetic inactivation, 10 flies were placed into arena chambers and placed in a 30°C incubator for 20 minutes prior to testing. The arena was then transferred to a preheated (30°C) light sealed box and connected to a humidified air delivery system. Flies were given an additional 15 minutes to acclimate to the box before recordings began. Group activity was recorded (33 frames/sec) for 20 minutes. Recorded .avi files of fly activity were processed by FFMPEG and saved as .mp4. Individual flies were tracked using Caltech Flytracker (Eyjolfsdottir et al., 2014) to obtain output features such as position, velocity, and angular velocity. Feature based activity was averaged across within each genotype and plots were generated in RStudio.

## Acknowledgements

K.M.S. and K.R.K were supported by the Smith Family Award Program for Excellence in Biomedical Research (K.R.K.) and the NIH grant R01AA024434 (K.R.K.). K.R.K. was also supported by a grant from NIH (5P20GM103645) to the Carney Institute for Brain Science Center for Nervous System Function COBRE. G.B., M.T., A.S. and J.D.F. were supported in part by NIH grants R01DC017146 and R01MH105368. We thank Vanessa Ruta for many fruitful discussions, advice on the manuscript and experiments, as well as support of R.C. We thank Janelia Fly facility for help with fly husbandry, and the FlyLight Project team for help with brain dissections, histological preparations, and confocal imaging performed at Janelia Research campus. We thank Hideo Otsuna and Takashi Kawaset (Janelia Research Campus) for helpful software advice and Arif Hamid (Brown University) for helpful programming advice. We also thank Gina Chieffallo for initial anatomical characterizations. This work was made possible in part by software funded by the NIH: Fluorender: An Imaging Tool for Visualization and Analysis of Confocal Data as Applied to Zebrafish Research, R01-GM098151-01.

## Notes

### Competing Interest Statement

The authors have declared no competing interest.

