## Supplementary Figures for "Transsynaptic mapping of *Drosophila* mushroom body output neurons"

[6] Current address: Howard Hughes Medical Institute, Department of Molecular and Cellular Biology, Harvard University, Cambridge, MA, 02138, USA

[7] Current address: Department of Biological Studies, Columbia University, New York, NY, 10027

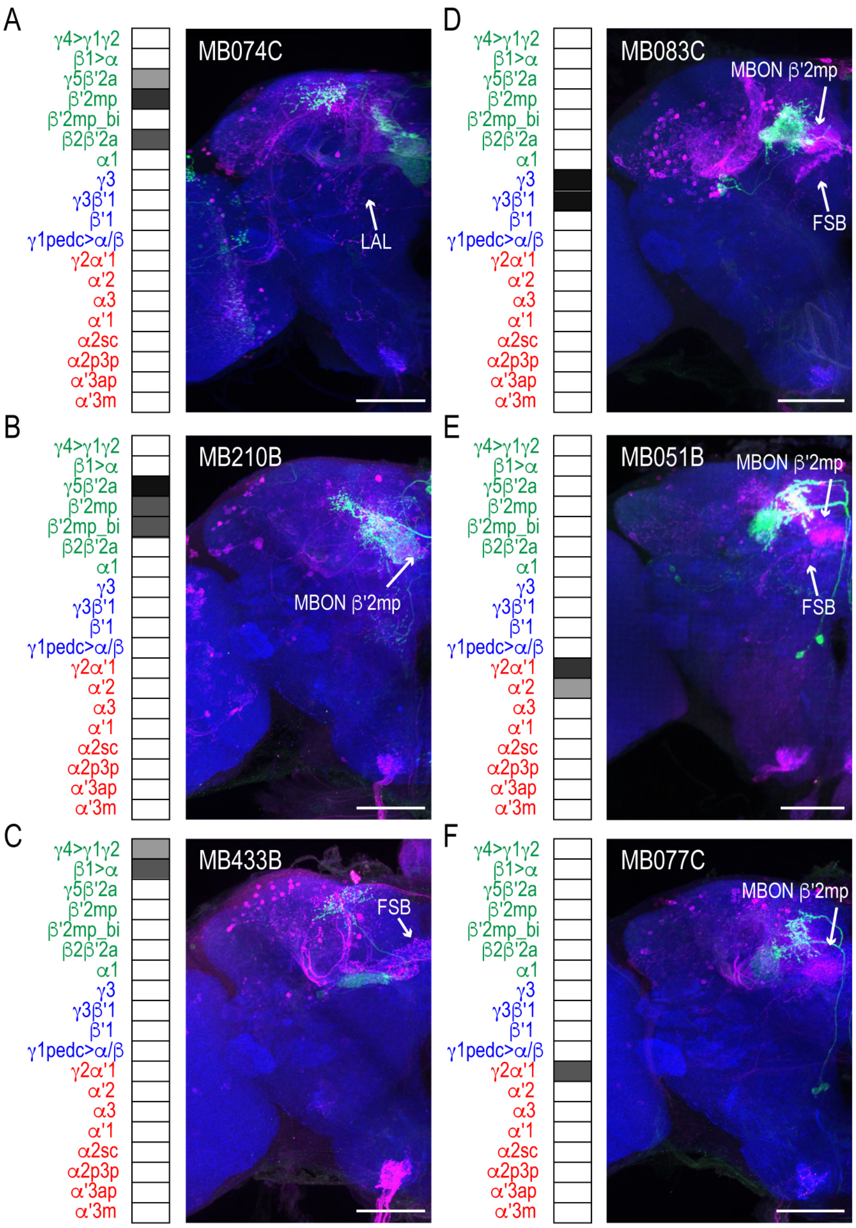

**Figure 1 Supplementary 1: MBON driver lines that have similar expression patterns also have similar postsynaptic connections across the brain.** Exemplar max-stacks of glutamatergic MBONs **(A)** MB074C, **(B)** MB210B, **(C)** MB433B, **(D)** GABAergic MBONs MB083C, and cholinergic MBONs **(E)** MB051B and **(F)** MB077C, *trans*-Tango identified postsynaptic connections. For max-stacks: green, presynaptic MBONs, magenta, postsynaptic *trans*-Tango signal, blue, *brp-SNAP* neuropil. A map of the MBONs that are included in the expression pattern in each driver line accompanies each exemplar with the relative expression pattern (greyscale, 1-5) accordingly to FlyLight (https://splitgal4.janelia.org/cgi-bin/splitgal4.cgi). MBON maps are organized by neurotransmitter type: green=glutamatergic, blue=GABAergic, red=cholinergic. Scale bar=50μm.

A

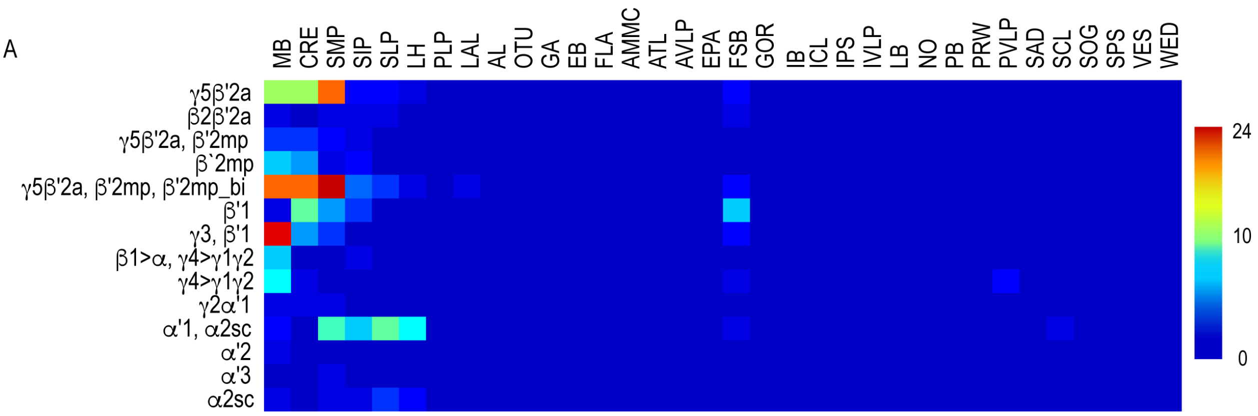

**Figure 2 Supplementary 1: Whole brain distribution of MBON postsynaptic connections overlap normalized across MBON driver lines. (A)** Heatmap displaying the overlap in signal of segmented MBON postsynaptic signal by brain region**.** Postsynaptic signal for each MBON was normalized across MBON driver lines allowing comparison of relative intensity of signal across the different MBON lines.

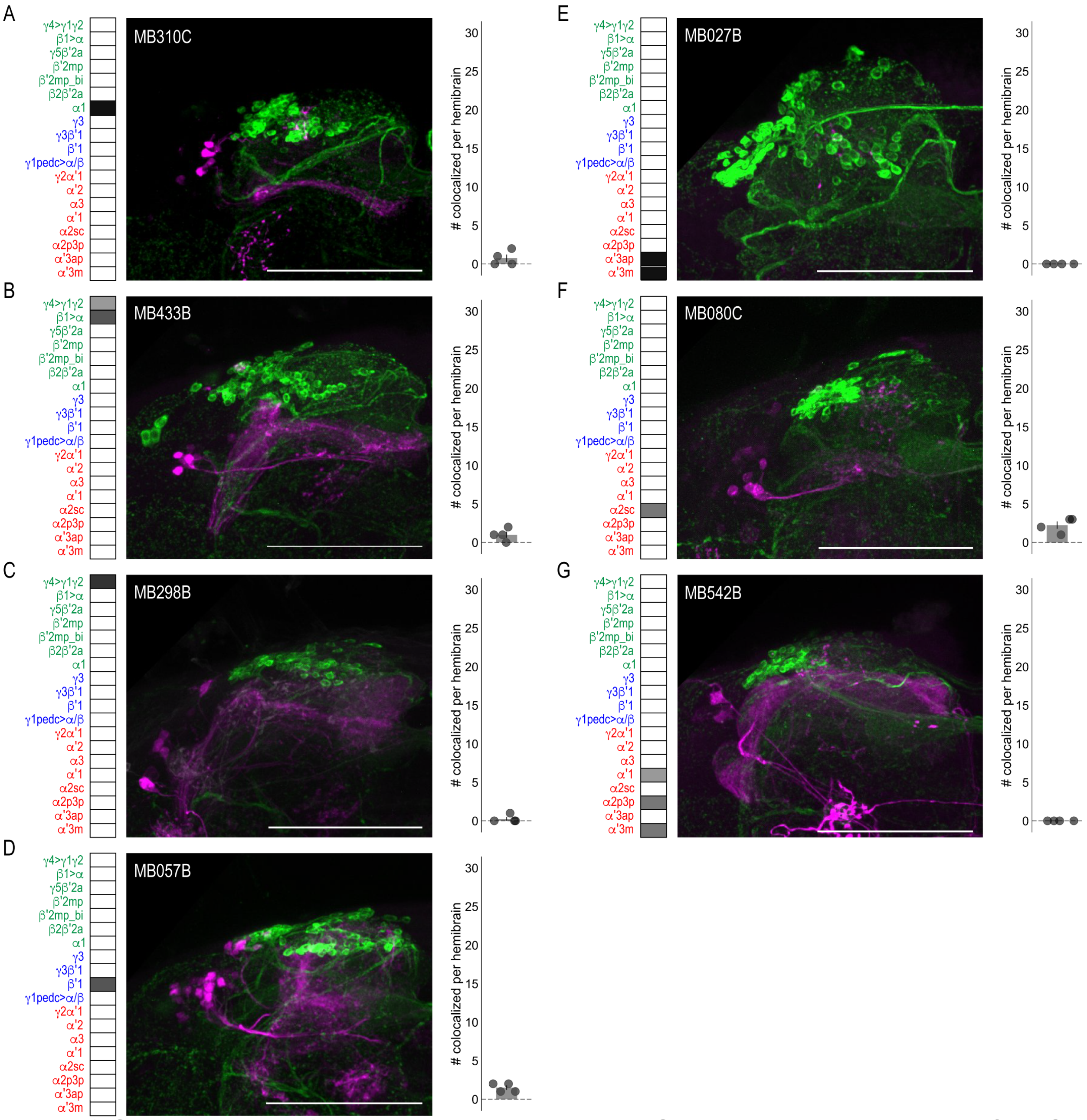

**Figure 3 Supplementary 1. DANs postsynaptic to MBONs.** Exemplar max-stacks of MBON lines in which TH+ cells were not overlapping or had few overlapping cells with postsynaptic signal of glutamatergic **(A)** MBON α1 (MB310C), **(B)** MBON γ4>γ1 γ2, β1>α (MB433B), **(C)** MBON γ4>γ1 γ2 (MB298B), **(D)** GABAergic MBON β′1 (MB057B), **(E)** cholinergic MBON α′3 (MB027B), **(F)** MBON α2sc (MB080C) and **(G)** MBONs α′1, α2p3p , α′3m (MB542B). Max stacks of MB are included, scale bar=50μm. Bar graphs indicate the average number of co-localized cells per hemisphere (mean +/- standard error). Green, TH-positive cells; magenta, postsynaptic *trans*-Tango signal. MBON maps are organized by neurotransmitter type: green=glutamatergic, blue=GABAergic, red=cholinergic.

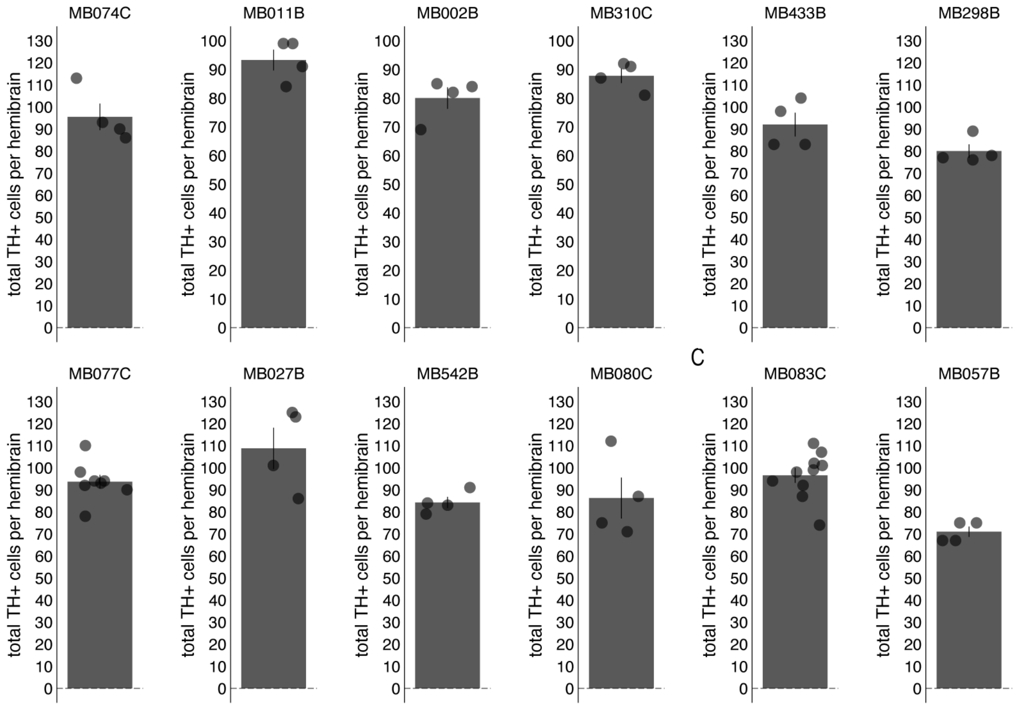

**Figure 3 Supplementary 2: Total PAM TH+ cells counted.** Bar graphs indicate the average number of PAM TH+ cells counted per hemisphere (mean +/- standard error).

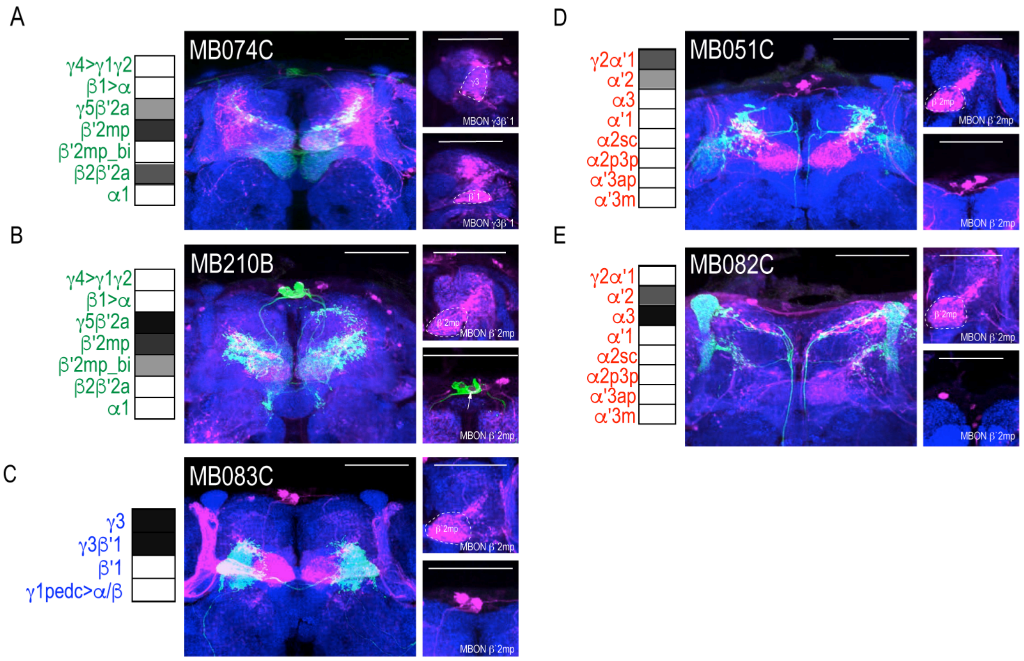

**Figure 4 Supplementary 1: Patterns of MBON convergence is consistent across MBON driver lines that have similar MBON expression**. MBON γ3β′1 receives input from **(A)** glutamatergic MBON β′2mp (MB074C). MBON β′2mp receives convergent input from **(B)** glutamatergic MBON γ5β′2a (MB210B), **(C)** GABAergic MBONs γ3, γ3β′1 (MB083C) and **(D)** cholinergic MBON γ2α′1 (MB051C) and **(E)** MBON α′2 (MB082C**).** For max-stacks: green, presynaptic MBONs, magenta, postsynaptic *trans*-Tango signal, blue, *brp-SNAP* neuropil, scale bar=50μm.

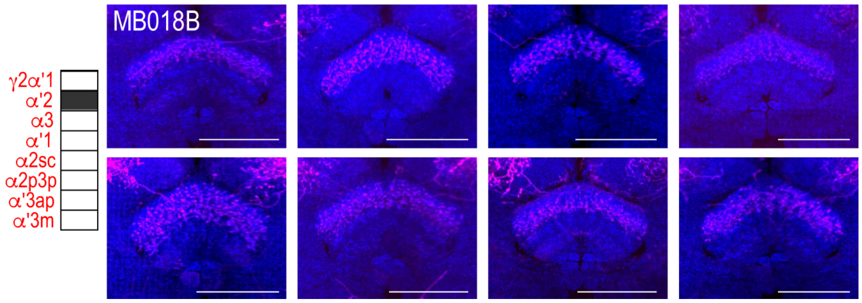

**Figure 5 Supplementary 1. Variability in FSB postsynaptic signal.** Exemplar max-stacks of cholinergic MBON α`2 whose postsynaptic neurons innervate the FSB highlighting the variability that existed in FSB innervation across eight different brains. Max-stacks are approximately 50μm thick. Magenta, postsynaptic *trans*-Tango signal; blue, neuropil. Scale bar=50 μm

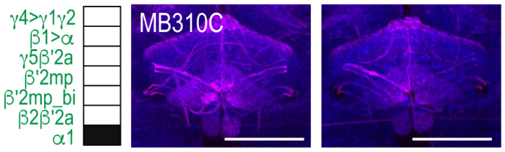

**Figure 5 Supplementary 2. MBON α1 postsynaptic signal innervating FSB in females.** Exemplar max-stacks of glutamatergic MBON α1 whose postsynaptic neurons innervate the FSB from two female flies highlighting the broad FSB innervation pattern of postsynaptic signa. Max-stacks are approximately 50μm thick. Magenta, postsynaptic *trans*-Tango signal; blue, neuropil. Scale bar=50 μm

A

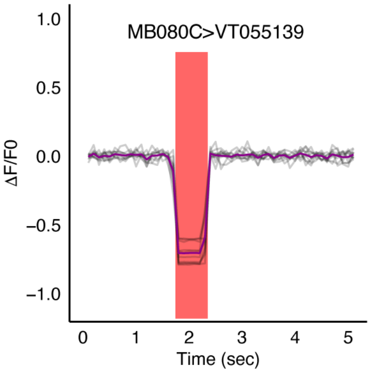

**Figure 7 Supplementary 1.** Optogenetic activation of MBON α2sc does not result in changes in signal recorded from of LAL neurons in SMP (VT015539).

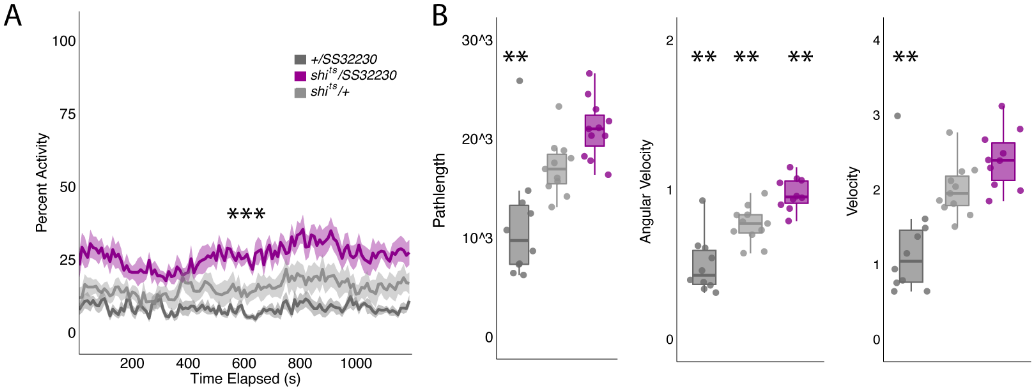

**Figure 7 Supplementary 2. (A)** *shibire*$^{ts}$ inactivation of LAL using split-GAL4 SS32230 resulted in significant increases in group activity (F(2,21)=13.94 p<0.0001). Group activity counts were binned over 10s periods, averaged across biological replicates of 10 flies each (n=8) and plotted against time. Lines depict mean +/- standard error. **(B)** One video was selected at random of each genotype and processed using FlyTracker to calculate the average pathlength (F(2,28)=14.76, p<0.0001), angular velocity (F(2,28)=28.16, p<0.0001) and velocity (F(2,28)=13.96, p<0.0001) of individual flies. Box plots with overlaid raw data were generated using RStudio. Each dot is a single fly. One-way ANOVA with Tukey Posthoc was used to compare mean and variance. ***p<0.0001, **p<0.001

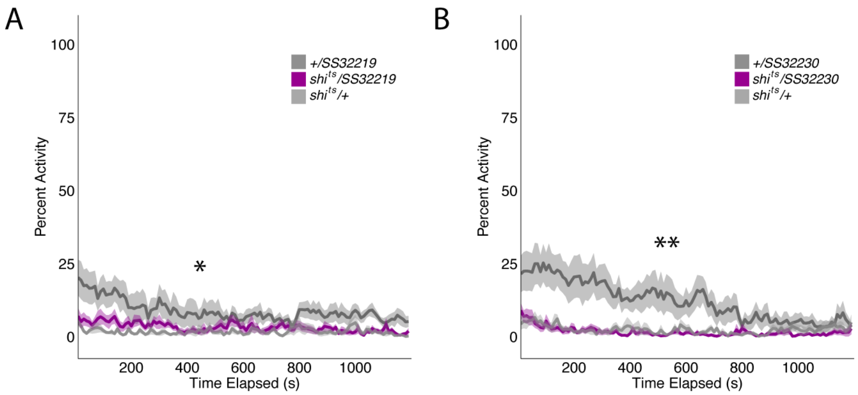

**Figure 7 Supplementary 3.** Group activity of split-GAL4 SS32219 and SS32230 at permission temperatures. Group activity of experimental groups were not significantly different from both genetic control groups. **(A)** Average group activity was significantly different at permissive temperatures for split-GAL4 line SS32219 ($F(2,21)=4.617$, $p=0.02$) although Tukey Post-hoc analysis revealed that *shi*$^{ts}$/SS32219 was not significantly different from either genetic control (SS32219/+ vs *shi*$^{ts}$/SS32219, p=0.08, *shi*$^{ts}$/+ vs *shi*$^{ts}$/SS32219, p=0.82). Instead, *shi*$^{ts}$/+ was significantly different from SS32219/+ (p=0.02). **(B)** Average group activity was significantly different at permissive temperatures for split-GAL4 line SS32230 ($F(2,21)=6.195$, $p=0.0078$) although Tukey Post-hoc analysis revealed that *shi*$^{ts}$/SS32230 was not significantly different from both genetic controls (SS32230/+ vs *shi*$^{ts}$/ SS32230, p=0.97, *shi*$^{ts}$/+ vs *shi*$^{ts}$/ SS32230, p=0.013). Instead, SS32230/+ was significantly different from both *shi*$^{ts}$/ SS32230 (p=0.013) and *shi*$^{ts}$/+ (p=0.02).
